# XomicsToModel: Omics data integration and generation of thermodynamically consistent metabolic models

**DOI:** 10.1101/2021.11.08.467803

**Authors:** German Preciat, Agnieszka B. Wegrzyn, Xi Luo, Ines Thiele, Thomas Hankemeier, Ronan M.T. Fleming

## Abstract

Constraint-based modelling can mechanistically simulate the behaviour of a biochemical system, permitting hypotheses generation, experimental design and interpretation of experimental data, with numerous applications, especially modelling of metabolism. Given a generic model, several methods have been developed to extract a context-specific, genome-scale metabolic model by incorporating information used to identify metabolic processes and gene activities in each context. However, existing model extraction algorithms are unable to ensure that a context-specific model is thermodynamically flux consistent. This protocol introduces `XomicsToModel`, a semiautomated pipeline that integrates bibliomic, transcriptomic, proteomic, and metabolomic data with a generic genome-scale metabolic reconstruction, or model, to extract a context-specific, genome-scale metabolic model that is stoichiometrically, thermodynamically and flux consistent. One of the key advantages of the `XomicsToModel` pipeline is its ability to seamlessly incorporate omics data into metabolic reconstructions, ensuring not only mechanistic accuracy but also physicochemical consistency. This functionality allows for more accurate metabolic simulations and predictions across different biological contexts, enhancing its utility in diverse research fields, including systems biology, drug development, and personalised medicine. The `XomicsToModel` pipeline is exemplified for extraction of a specific metabolic model from a generic metabolic model; it enables omics data integration and extraction of physicochemically consistent mechanistic models from any generic biochemical network. It can be implemented by anyone who has basic MATLAB programming skills and the fundamentals of constraint-based modelling.

**Key points:** XomicsToModel is a semi-automated pipeline that integrates bibliomic, transcriptomic, proteomic, and metabolomic data with a generic genome-scale metabolic reconstruction or model.

It enables the seamless incorporation of multi-omics datasets into metabolic reconstructions, ensuring mechanistic accuracy and physicochemical consistency.

## Introduction

### Genome-scale metabolic models

Genome-scale metabolic models are computational frameworks that simulate the metabolic processes within an organism. They are essential tools for understanding complex biochemical networks and predicting physiochemical and biochemically realistic metabolic fluxes. The main goal of a genome-scale metabolic model is to represent known metabolic functions in living systems to facilitate hypotheses generation, experimental design, and interpretation of experimental data [1].

Genome-scale metabolic models can be generic, covering multiple tissues, organisms or organelles, or they can be specific, tailored to specific tissues, cell types, or environmental conditions. There are four complementary approaches for the development of a genome-scale metabolic models:

1. From scratch: If no network reconstruction is available, a high-quality genome-scale metabolic reconstruction of a single organism can be generated from scratch by following established protocols [2]. Once this reconstruction is in place, it can be used to derive various metabolic models by computational application of different combinations of mathematical modelling assumptions.
2. Generic reaction databases: Given a generic metabolic reaction database, containing metabolic reactions from multiple organisms, an organism-specific model can be derived directly from this data. There are a several existing methods for doing this, e.g., CarveMe [3].
3. Orthologous genes: If a network reconstruction, or a model, is available for an organism with orthologous genes, it is possible to generate metabolic models for related species. Methods for microbial [4] and mammalian [5] species have been proposed.
4. Tailoring models: Given a generic metabolic reconstruction or model for a multi-cellular organism (containing metabolic reactions from any anatomical region), there are several methods that can be used to derive a model that is specific for a particular anatomical region [6].

In all except the first instance, the experiment starts with an existing generic reconstruction or model and the method extracts a specific model, which is a subset of the generic reconstruction.

A specific model may be specific to a particular organism or a particular anatomical region. For example, given a generic human metabolic model, one could extract a hepatocyte model [7], with the capability to model different contexts, depending on the constraints subsequently applied to it. A specific model may also be context-specific, which is a particular organism or anatomical region in a particular environmental and internal context. From the same starting point, with different input data, one could extract a context-specific model, e.g., a hepatocyte model that is specific to a fasting state [8]. Context-specific data can include both experimental results and bibliomic data. When discrepancies arise, such as genes, metabolites, or reactions present in one data source but absent in another, preference must be given to either experimental or bibliomic data, resulting in two types of models: cell-type-specific and condition-specific. Cell-type-specific models prioritise experimental data, while condition-specific models prioritise bibliomic data [9, 10].

To ensure the accuracy and biological relevance of a context-specific model, it is crucial to establish stoichiometric, flux, and thermodynamic consistency. Stoichiometric consistency guarantees that all reactions within the model are mass-balanced, ensuring that the inputs and outputs of each reaction align with the conservation of mass [11]. Flux consistency ensures that every reaction in the model can carry a non-zero steady-state flux, allowing for meaningful metabolic simulations [12]. Lastly, thermodynamic flux consistency ensures that the model adheres to the laws of thermodynamics, meaning that each reaction can carry a non-zero net flux in a thermodynamically feasible direction, respecting energy conservation and equilibrium constraints [13]. These consistency checks are essential to generate a physiologically plausible and predictive metabolic model.

Established model extraction methods ensure that a context-specific model is flux consistent. This concept is explained mathematically after a brief definition of the mathematical notation used throughout.

- *R* denotes the field of real numbers
- *R*_*n*_ denotes the vector space of *n*-tuples of real numbers
- *R*^*m*×*n*^ denotes the space of *m* × *n* matrices with entries in *R*
- Greek lowercase and uppercase letters denote scalars and sets, respectively.
- Roman lowercase and uppercase letters denote vectors and matrices, respectively.
- *N*^*T*^denotes the transpose of a matrix *N*.
- ‖*r*‖ denotes the zero norm of the vector *r*.

In any optimisation problem, the variables are indicated below the first symbol in the optimisation problem definition, e.g., denotes maximisation of an objective in an optimisation problem where the only variable is *v* and all other symbols are input data to the optimisation problem.

To predict the steady-state behaviour of metabolic networks, Flux Balance Analysis is commonly employed as it provides an efficient and widely recognised method for calculating metabolic fluxes under given constraints. Flux Balance Analysis ensures that the metabolic model adheres to mass balance and other physiological constraints, making it a foundational approach for exploring metabolic capabilities. This method serves as a stepping stone for more advanced flux analysis approaches, which may offer additional insights or higher complexity, but FBA remains essential for establishing baseline flux distributions.

#### Flux Balance Analysis

Flux Balance Analysis [14] requires the solution to the following optimisation problem:

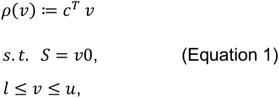

where *v* ∈ *R*^*r*^ is the net rate of each reaction, and *ρ*(*v*) ≔ *c*^*T*^*v* is a biologically motivated linear objective, specified by the coefficient vector *c* ∈ *R*^*r*^. The matrix *S* ∈ *R*^*m*×*r*^ is a stoichiometric matrix, where *m* is the number of metabolites, and *r* is the number of internal plus exchange reactions. The constraint *Sv* = 0 represents the assumption of steady-state, that is:

- Production = consumption for metabolites not exchanged across the boundary of the system.
- Production + uptake = consumption + secretion for metabolites exchanged across the boundary of the system.

The inequalities *l* ≤ *v* ≤ *u* denote box constraints from the lower (*l* ∈ *R*^*r*^) and upper bounds (*u* ∈ *R*^*r*^) on reaction rates.

A reaction is considered flux consistent if it can carry a non-zero steady-state flux, meaning there is at least one steady-state flux vector *v* in the feasible set *Ω* ≔ {*Sv* = 0, *l* ≤ *v* ≤ *u*} where *v*_j_ ≠ 0. This implies the reaction can function under the given conditions and constraints. Conversely, a reaction is considered flux inconsistent if its rate *v*_j_ ≠ 0 for all steady state flux vectors in the feasible set *Ω*. Since Flux Balance Analysis only predicts steady-state flux, including flux inconsistent reactions in a model can be misleading. Therefore, it is important to distinguish between:

- Reactions that cannot carry steady state flux (independent of the objective chosen)
- Reactions that do not carry steady state flux in a particular Flux Balance Analysis solution (dependent of the objective chosen).

Therefore, flux inconsistent reactions should be identified in a model before interpreting the results of methods such as Flux Balance Analysis. In a metabolic reconstruction, a flux inconsistent reaction (supported by experimental evidence) may serve as a starting point for refinement (e.g., by gap filling [15]) that adds one or more additional reactions to a network to enable flux through a formerly flux inconsistent reaction.

#### Thermodynamic feasibility and thermodynamic flux consistency

It is well established that Flux Balance Analysis requires additional constraints to ensure that a steady state flux vector is also thermodynamically feasible [16, 17], i.e., consistent with energy conservation and the second law of thermodynamics [18]. The critical necessity at genome-scale is to incorporate any additional constraints (or terms in the objective) in a mathematical form that, when computationally implemented, retains the computational efficiency and certificates available with linear optimisation. As exemplified in Figure 1, a stoichiometric matrix may be split into two sets of columns *S* ≔ [*N, B*], where the matrix *N* ∈ *R*^*m*×*n*^ represents internal reactions, which do conserve mass, and the matrix *B* ∈ *R*^*m*×*k*^ represents external reactions, which do not conserve mass, e.g., the reaction with equation *A* → ∅. Within this framework, chemical reactions that do not conserve mass serve as modelling constructs for simulating the exchange, demand, or sink of metabolites with the environment. Exchange reactions model the flux of metabolites into or out of the system, simulating substance exchange with the external environment. Demand reactions represent the cellular need for a specific metabolite, mimicking the consumption or requirement within the biological system. Lastly, sink reactions act as outlets for metabolites, replicating their removal from the system, whether through excretion, degradation, or otherwise removed from the cellular environment.

**Figure 1:**
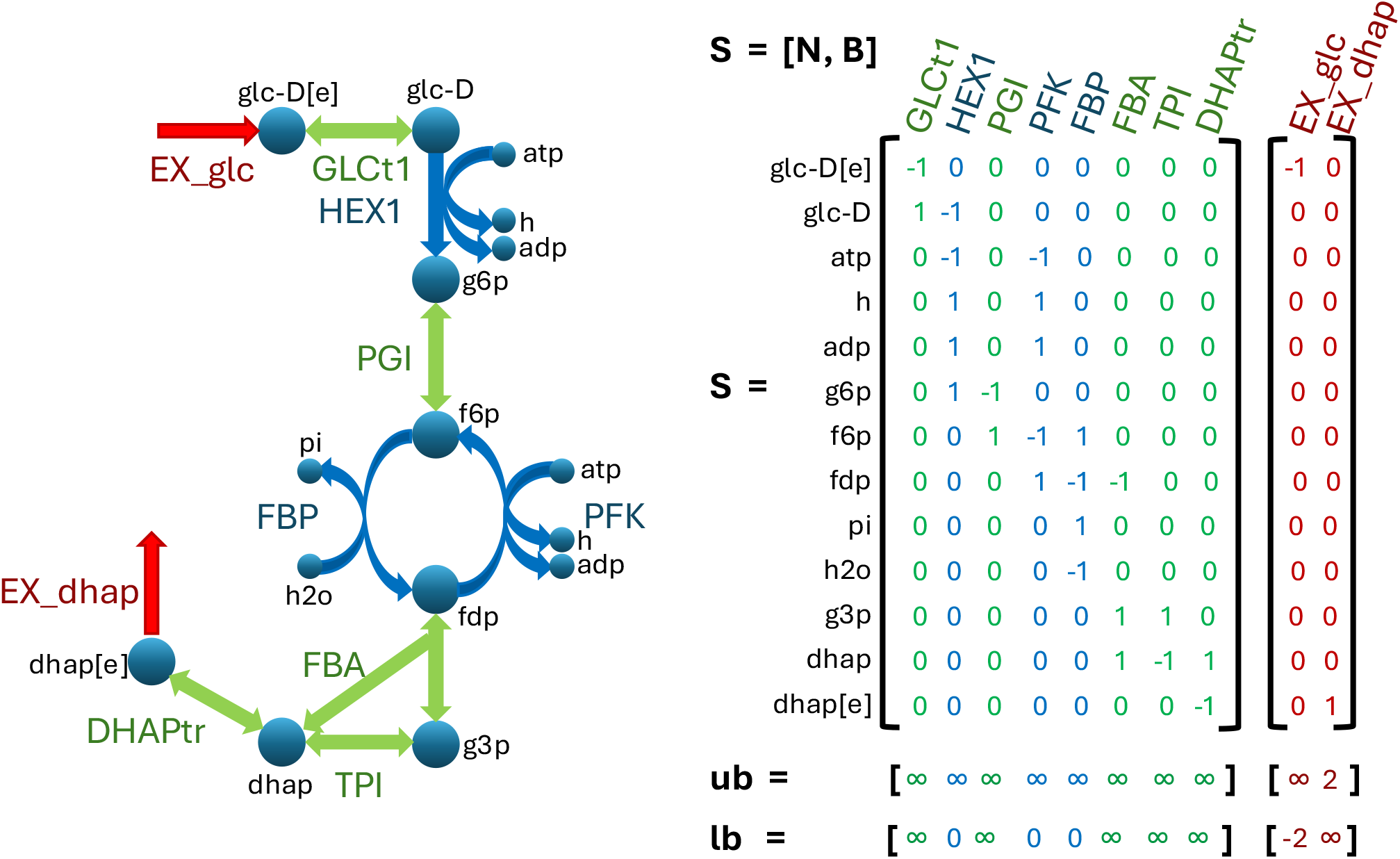
Network diagram of upper glycolysis (left) with reactions (arrows, upper case labels) and metabolites (nodes, lower case labels) with corresponding labels. The corresponding Stoichiometric matrix (right, *S* ∈ *R*^13×10^, *N* ∈ *R*^13×8^, *B* ∈ *R*^13×2^), with reversible reactions shown in green, non-reversible reactions shown in blue, and exchange reactions shown in red. The upper bound (*u*) of internal reactions is unconstrained, whereas the lower bound (*l*) is limited by reaction directionality or by the maximum uptake rate, which can be seen as a constraint on uptake of extracellular glucose (glc_D[e]) or secretion of dihydroxyacetone phosphate (dhap[e]) from the environment. All metabolite and reaction abbreviations are with respect to the namespace in www.vmh.life [19].

By splitting internal net fluxes into unidirectional fluxes, and maximising the entropy of unidirectional fluxes, one can compute a thermodynamically feasible steady state net flux vector in a genome-scale biochemical model [13]. Maximising the entropy of unidirectional fluxes is a convex optimisation problem that retains the efficiency as well as feasibility certificates available with linear optimisation [20]. This approach of maximising the entropy of unidirectional fluxes assumes that the model constraints admit a solution that corresponds to a thermodynamically feasible net flux vector. However, we define a model to be thermodynamically flux consistent if each of its reactions admits a nonzero net flux in at least one thermodynamically feasible, steady-state, net flux vector. Chemically, a thermodynamically feasible net flux vector may be zero for a reaction if the substrates and products of a reaction are in thermodynamic equilibrium. Biochemically, a thermodynamically feasible net flux vector may be zero for a reaction (effectively zero at a biochemically relevant timescale) if the enzyme corresponding to that reaction is absent, even if the substrates and products for that reaction are not in thermodynamic equilibrium. We define a reaction to be thermodynamically flux consistent if it admits a non-zero net flux that is also thermodynamically feasible to eliminate thermodynamically inconsistent reactions that only admit a zero net flux in a thermodynamically feasible flux vector. In summary, a reaction is flux consistent if it admits a nonzero flux that satisfies mass balance and bound constraints, while it is thermodynamically flux consistent if it admits a non-zero net flux that satisfies a combination of mass balance, bound constraints, energy conservation and the second law of thermodynamics.

A thermodynamically feasible flux does not exist when at least one combination of bounds on reaction rates forces net flux around a stoichiometrically balanced cycle [21]. Consider the following stoichiometrically balanced cycle of reactions *A* ⇌ *B* ⇌ *C* ⇌ *A*. If the bounds are such that *A* → *B* → *C* → *A* is the only feasible net flux, then the network containing this cycle cannot carry a thermodynamically feasible steady state net flux. The second law of thermodynamics requires that a chemical potential difference between substrates and products is required for net flux. Let *μ*_*a*_ denote the chemical potential of metabolite *A*. Given net flux *A* → *B*, the second law of thermodynamics requires *μ*_*A*_ > *μ*_*B*_ and similarly *μ*_*B*_ > *μ*_*C*_ and *μ*_*C*_ > *μ*_*A*_. However, the first pair of inequalities imply *μ*_*A*_ > *μ*_*C*_, which is inconsistent with the last, unless one assigns more than one chemical potential to *C* at the same instant. However, to do so would be inconsistent with energy conservation, which requires each row of a stoichiometric matrix corresponds with a single chemical potential, assuming each row represents a metabolite in a specific compartment [18].

Stoichiometrically balanced cycles are biochemically faithful network topological features that are omnipresent in genome-scale models. It is not the presence of a stoichiometrically balanced cycle, per se, that presents a problem. Thermodynamically inconsistent specification of reaction bounds may force net flux around a stoichiometrically balanced cycle and thereby prevent the prediction of a thermodynamically feasible net flux. Also, failure to implement thermodynamic constraints on the optimal solution of an optimisation problem may lead to the prediction of a thermodynamically infeasible net flux. As recognised by Desouki et al. [21], provided that a model does not contain a combination of thermodynamically inconsistent bounds, minimisation of the one-norm of net flux, subject to additional constraints maintaining the direction of net flux, is guaranteed to remove that part of a steady state flux vector that is thermodynamically infeasible. This approach is attractive, as it is based on linear optimisation, but it has yet to be leveraged for generation of thermodynamically flux consistent models. Generation of thermodynamically flux consistent models should, in principle, increase the biochemical fidelity of constraint-based models as it opens up the possibility of leveraging established methods to efficiently compute thermodynamically feasible steady state fluxes [13, 21] that assume an input model is thermodynamically flux consistent.

Given a generic reconstruction or model, all established model extraction algorithms, including GIMME [22], iMAT [23], MBA [7], mCADRE [24], FastCore [12], and CarveMe [3], as well as gap-filling algorithms [15], extract a specific model based on either a binary (present/absent) or weighted assignment of reactions desired to be present in a specific model, where each reaction is (net) flux consistent. However, these algorithms do not ensure that the resulting models are thermodynamically flux consistent. This protocol introduces an enhanced model extraction algorithm (`thermoKernel` algorithm within a model generation pipeline that generates models that are thermodynamically flux consistent.

### Development of the protocol

This protocol presents the `XomicsToModel` pipeline, which enables extraction of a context-specific, genome-scale metabolic model that is stoichiometrically, thermodynamically and flux consistent from generic reconstruction and omics data. The pipeline was initially developed and validated in the context of dopaminergic neuronal metabolism [9], where transcriptomic, metabolomic and bibliomic data were integrated with Recon3D [25]. Beyond this first application, `XomicsToModel` has already been successfully used in several peer-reviewed studies. For example, it has been applied to construct context-specific genome-scale models of human dopaminergic neurons by integrating transcriptomic and exometabolomic data to investigate neuronal metabolism and its bioenergetic vulnerabilities [26]; to integrate multi-omics datasets for the derivation of condition-specific human metabolic models, demonstrating reproducibility across biomedical contexts [27]; and to explore disease mechanisms by combining transcriptomic, proteomic, and metabolomic data [28]. In addition, XomicsToModel has been applied to model macrophage metabolism in Gaucher disease, where simulations predicted mitochondrial impairment, a glycolytic shift, and dysregulation of cholesterol and sphingolipid metabolism—offering mechanistic insight into lysosomal dysfunction and potential therapeutic targets [29]. In parallel, the pipeline has also been employed to predict fluxes when combined with stable isotope labelling data [30, 31]. Together, these applications demonstrate the robustness and adaptability of the pipeline across neuroscience and broader biomedical research.

Developing a comprehensive pipeline for generating thermodynamically flux consistent models is essential for enhancing the accuracy of constraint-based metabolic models. By incorporating methods like the `thermoKernel` algorithm, this pipeline addresses the limitations of existing model extraction algorithms and ensures that thermodynamic consistency is integral to model generation. This holistic approach enables more reliable predictions of metabolic behaviour and flux distributions, ultimately advancing our understanding of cellular processes.

### Overview of the procedure

A metabolic reconstruction is a representation of a metabolic network whose content may not entirely satisfy modelling assumptions. By contrast, a metabolic model is a mathematically defined representation that adheres to such assumptions. `XomicsToModel` is a semi-automated computational pipeline designed to generate context-specific metabolic models from a generic reconstruction or model, combined with omics data and user-defined technical parameters.

The pipeline consists of 23 sequential steps (Fig. 2), organized into four functional stages: (i) input preparation, in which omics datasets, technical parameters, and reconstruction metadata are specified; (ii) data integration and consistency checks, where context-specific information is mapped onto the generic reconstruction; (iii) model tailoring, in which constraints are iteratively applied and internal variables refined; and (iv) model finalization, where a comprehensive record of the reconstruction process is generated, including the reactions added or removed at each step. Each step is controlled by user-defined variables, and, except for the final stage, produces intermediate outputs that guide iterative refinement.

**Figure 2:**
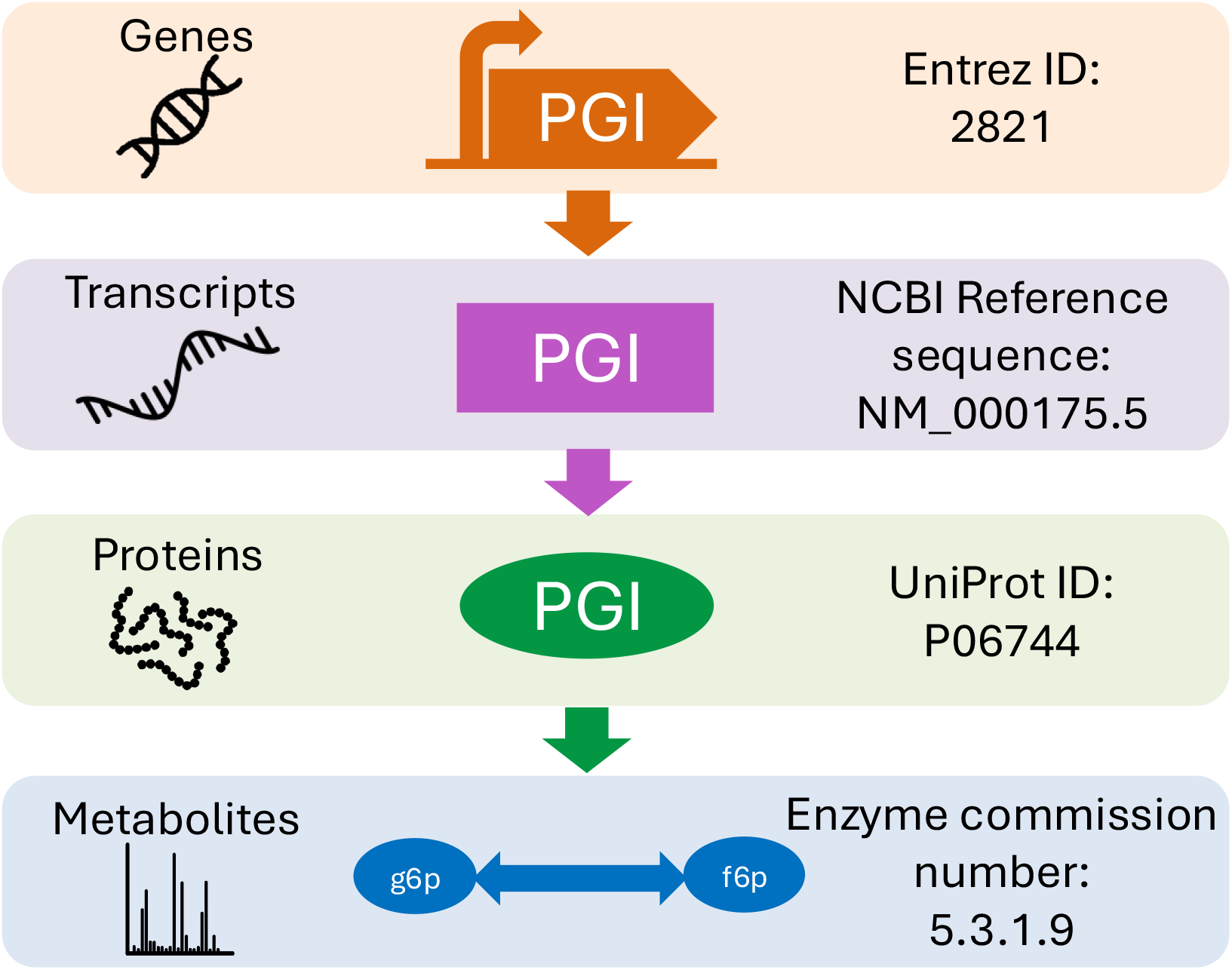
Overview of the `XomicsToModel` pipeline. It is possible to create genome-scale models iteratively and systematically using the `XomicsToModel` pipeline. The generated models can be used to design or interpret the information based on bibliomic data from manual literature curation (red), as well as metabolomic (blue), proteomic (green) or transcriptomic (purple) data. Furthermore, experimental data can be used to validate or refine the context-specific model. The `XomicsToModel` pipeline (grey) extracts a new model by integrating a generic model with context-specific information based on the technical parameters such as the extraction algorithm, method for identifying active genes, or maximum reaction rates. Steps 12, 13, and 22 are indented as they represent subprocesses within the pipeline.

Although a draft model can be generated using default parameters, optimal results require adjusting parameters to the biological context and validating model predictions against experimental data. A practical strategy is to generate an ensemble of draft models under different parameterizations, quantify their predictive performance, and select the context-specific model that best aligns with experimental evidence [9].

For clarity, the full sequence of the 23 modular steps is provided below, ensuring reproducibility and highlighting the adaptability of the pipeline to available data.

The workflow of `XomicsToModel` proceeds in a sequence of modular steps, ensuring that the extraction can adapt to the available information. It begins with data preparation (Step 1) followed by a generic model check (Step 2) to assess consistency, after which an objective function can be set if desired (Step 3). The model can then be enriched with missing reactions (Step 4), core metabolites and reactions are identified (Step 5), and reaction bounds established (Step 6). Gene activity is incorporated (Step 7), with optional closure of ionic fluxes (Step 8), exchange reactions (Step 9), and sink or demand reactions (Step 10). At this stage, metabolic constraints can be introduced (Step 11), drawing from cell culture data (Step 12) and metabolomics (Step 13), while allowing custom user-defined constraints (Step 14) or coupled reactions (Step 15). The model is refined by removing inactive reactions (Step 16) or genes (Step 17), and subsets that are flux-consistent (Step 18) or thermodynamically consistent (Step 19) are identified if required. Active reactions based on gene expression are then integrated (Step 20), before the final extraction of the context-specific model (Step 21), which can also be performed using `thermoKernel` if selected (Step 22). Lastly, the workflow completes with final adjustments and reporting (Step 23). Importantly, the process is modular: if transcriptomic, metabolomic or bibliomic data are missing, the extraction still proceeds seamlessly with the available information.

#### Input data

The input data to the `XomicsToModel` pipeline is a generic biochemical network and a set of omics data. The generic biochemical network may be a reconstruction or a model without any specificity, and is not required to be stoichiometrically, thermodynamically or flux consistent. The `XomicsToModel` pipeline allows a flexible and modular integration of transcriptomic, proteomic, and metabolomic data, as well as bibliomic data abstracted from literature curation. In each case, the input omics data may be qualitative (present, absent, unspecified), semiquantiative, quantitative, or combinations thereof. The `XomicsToModel` pipeline is complemented by functions to automatically import omics data. In the `XomicsToModel` pipeline, metabolomic data on intracellular metabolites can currently only be integrated in a qualitative or semi-quantitative, rather than fully quantitative manner. As `XomicsToModel` enables generation of a stoichiometrically, thermodynamically, and flux consistent model, this model is compatible with subsequent integration of quantitative intracellular metabolomic data using a variety of established techniques [32].

The application of the `XomicsToModel` pipeline to extract a dopaminergic neuronal metabolic model [9] demonstrates its flexibility with respect to incorporation of a variety of qualitative and quantitative constraints. For example, information about the presence or absence of metabolites in the culture medium, together with quantitative metabolite exchange reaction rates, was integrated into the model. These exchange rates were used to set bounds on the corresponding reactions through quadratic optimization, with each constraint weighted by the inverse of its measurement uncertainty. In the `XomicsToModel` pipeline, it is possible to manually specify metabolic reactions to extend an input reconstruction or model (with manually specified metabolic reactions) when a generic model is missing certain key pathways relevant for a system of interest.

#### Constraint relaxation

The `XomicsToModel` pipeline incorporates a series of tests for flux and optionally thermodynamic consistency after each step, e.g., after removal of generic metabolites or reactions assigned not to be present in a specific model. This approach ensures the early detection of infeasible constraints resulting from inaccurate or inconsistent experimental data. Detection of inconsistency is followed by algorithmic relaxation of constraints (e.g. bounds) to render the draft model feasible. Specifically, the `XomicsToModel` pipeline automatically searches for the minimal number of constraint relaxations required to admit a flux consistent model in the case that inconsistent or incorrect omics data renders a draft model infeasible. This approach can be used to feed back into data processing or input to avoid model infeasibility, which is often a challenge with omics data integration. For example, relaxation of a steady state constraint points to the need for additional reactions to enable either synthesis or degradation of a metabolite where the existing total rate of synthesis (production) cannot match the total rate of degradation (consumption). In this case, additional reactions may need to be added to the generic model based on following established manual reconstruction protocols [2], or by recourse to algorithmic gap filling [15].

The application of the `XomicsToModel` pipeline to extract a dopaminergic neuronal metabolic model [9] presented a novel challenge for constraint-based modelling because the extracted model was required to be representative of neurons that do not grow. Therefore, the maximisation of biomass growth could not be used as an objective. Instead, a set of coupling constraints [35] were added to represent cell maintenance requirements, e.g., for the turnover of key metabolites. However, if incorrectly scaled with respect to constraints on exchange reactions, the application of coupling constraints may generate an infeasible draft model. Therefore, an algorithmic proposition of a minimal number of constraint relaxations accelerates the identification of biochemically inconsistent constraints and the refinement of input data. That is, proposed relaxations should be reviewed manually for consistency with additional biochemical knowledge then adjustments made to ensure that the input data, the input generic model, or both, are consistent with a feasible model.

#### Model extraction

The `XomicsToModel` pipeline is compatible with various model extraction algorithms (Table 1), with an established interface to FastCore [12]. In addition, the `XomicsToModel` pipeline builds upon `thermoKernel`, a novel algorithm used to extract the dopaminergic neuronal metabolic model [9], but applicable for the extraction of any context-specific model that is required to be stoichiometrically, thermodynamically and flux consistent. The ability of `thermoKernel` to ensure thermodynamic flux consistency during the model extraction process opens new possibilities for data integration. Thermodynamic consistency is implemented by constraining the possible relationships between reaction flux and metabolite chemical potential. Consequently, it is possible to directly specify a qualitative (presence, absence, unspecified) or quantitative (weighted) assignment of metabolites desired to be present in a specific model.

**Table 1:** Comparison of technical features of algorithms for extraction of a context-specific model, given a generic model of a single organism.

| Properties \ Algorithms | GIMME | iMAT | MBA | tINIT | mCADRE | FastCore | thermoKernel |
| --- | --- | --- | --- | --- | --- | --- | --- |
| Citation | [22] | [23] | [7] | [34] | [24] | [12] | N/A |
| Year published | 2008 | 2010 | 2010 | 2014 | 2012 | 2014 | 2023 |
| Metabolite list | × | × | × | × | × | × | ✓ |
| Metabolite weights $\in R$ | × | × | × | × | ✓ | × | ✓ |
| Active reaction list | ✓ | ✓ | ✓ | ✓ | ✓ | ✓ | ✓ |
| Reaction weights $\in R$ | × | × | ✓ | ✓ | ✓ | × | ✓ |
| Coupling constraints | × | × | × | × | × | ✓ * | ✓ |
| Thermodynamic consistency | × | × | × | × | × | × | ✓ |
| Constraint relaxation | × | × | × | × | × | × | ✓ |
| Algorithmic approach | MILP | MILP | FVA | MILP | fastFVA | QC-LP | DCA-LP |
| MILP, mixed integer linear programming; FVA, flux variability analysis, fastFVA [33], computational acceleration of flux variability analysis; QC-LP, quasiconcave sequence of linear programs; DCA-LP, difference of convex function sequence of linear programs. * denotes subsequently implemented in the COBRA Toolbox [38]. In a context-specific metabolic network, a metabolite is considered to be present if the sum of its rate of production plus consumption is above a minimum magnitude considered non-zero (typically 1e-4), otherwise it is considered to be absent. |  |  |  |  |  |  |  |

More generally, it is possible to specify any combination of qualitative assignments and quantitative weights to metabolites and reactions to reflect what is desired to be present or absent from a specific model. These weights are determined objectively based on omics evidence to avoid any subjective errors in assignment. This enables the `thermoKernel` algorithm to extract a thermodynamically flux consistent model that is a trade-off between qualitative assignments and/or quantitative weights of rows and columns of a stoichiometric matrix. The application of the `XomicsToModel` pipeline to extract a dopaminergic neuronal metabolic model [9] demonstrates this flexibility with qualitative and weighted specification of metabolites and reactions desired to be present in the dopaminergic neuronal model. For example, the `XomicsToModel` pipeline enables integration of quantitative weights on reactions from transcriptomic data at the same time as qualitative assignments from metabolomic data, where the metabolite presence (absence or unknown) can now be applied directly as context-specific data input.

#### Ensemble modelling

The `XomicsToModel` pipeline is flexible, and various additional functions may be incorporated into it. The supplementary `XomicsToMultipleModels` function can be used to generate an ensemble of context-specific genome-scale models by varying the specific data and technical parameters, as well as experimental data or generic models used in the model extraction process. Moreover, a supplementary `modelPredictiveCapacity` function can be added to estimate the predictive capacity of an extracted model, given independent data. The creation of multiple models with varying parameters and comparison of the predictive capacity of each model, enables identification of optimal input data and technical parameters. For example, where multiple different data types are available to generate a context-specific model (e.g., transcriptomic data corresponding to genes that encode metabolic enzymes, information manually curated from the literature on the presence or absence of activity on particular metabolic reactions), it is not always obvious what the relative merits of those input data are, especially as they may be conflicting. For example, gene expression might indicate that the gene for an enzyme is expressed, but manual curation indicates that the corresponding reaction is not active. In this situation, a pragmatic approach is to generate an ensemble of models, then compare their predictive capacity, and then select the combination of input data and technical parameters that provides the most accurate prediction. Further information on ensemble modelling may be found in the published literature [36].

### Applications

One of the first applications of `XomicsToModel` was to construct dopaminergic neuronal models, providing insight into metabolic dysfunction in Parkinson’s disease [9]. Substantia nigra dopaminergic neurons are the most susceptible to degeneration in Parkinson’s disease, yet its biochemical mechanisms remain poorly understood [37]. This foundational work has been extended in other biomedical contexts. For example, `XomicsToModel` was employed in studies of human neuronal metabolism [26], in condition-specific human metabolic reconstructions integrating diverse omics datasets [27], and in disease-focused analyses where transcriptomic, proteomic, and metabolomic integration revealed mechanistic alterations [28]. Its application to Gaucher disease macrophages [29] showed how the protocol can reveal systemic metabolic reprogramming, identifying mitochondrial dysfunction, altered lipid metabolism, and transcriptional regulators as drivers of disease phenotypes. In addition, ongoing studies have successfully used the pipeline to integrate isotopic labelling experiments with constraint-based modelling [30, 31].

This approach, guided by the `XomicsToModel` pipeline, offers broader applications for investigating metabolic dysfunction in various diseases. More generally, the `XomicsToModel` pipeline can be applied to any situation where one has a generic (or universal) reconstruction (or model) and seeks to extract a specific subset of it, based on transcriptomic, proteomic, or metabolomic data, or combinations thereof. This pipeline does not impose any requirements regarding the species identity in the input reconstruction or model, nor does it limit the number of species represented in either the input or output models. The most computationally intensive steps are the identification of the largest thermodynamically consistent subset of an input reconstruction or model and the extraction of a thermodynamically consistent subset of minimal size, both of which are achieved with the `thermoKernel` algorithm. As the `thermoKernel` algorithm is based on a sequence of linear optimisation problems, its performance scales accordingly. Qualitative specification of metabolite (presence/absence/unspecified), reaction (activity/inactivity/unspecified), gene (activity/inactivity/unspecified), and combinations thereof are all possible. Weighted specification to bias for or against the inclusion of metabolites, reactions, or both (based on input transcriptomic, proteomic, or metabolomic data) can be performed alone or in combination with a qualitative specification. The `XomicsToModel` pipeline is interfaced via a function, but multiple options can be specified to enable modular use of the `XomicsToModel` pipeline to implement model extraction priorities reflective of a wide variety of scenarios. As such, the `XomicsToModel` pipeline will likely have widespread applicability.

### Comparison with other methods

The `XomicsToModel` pipeline is integrated within the COBRA Toolbox [38]. It consists of a suite of functions for context-specific input data preprocessing, identification of the subset of a reconstruction that satisfies certain modelling assumptions, integration of constraints derived from context-specific data with a generic metabolic model, recovery of model feasibility if the constraints are inconsistent, extraction of a context-specific model, or an ensemble thereof. Besides, it contains a wide variety of functions for exploring the properties of generated models, including evaluation of their predictive fidelity, as detailed in the Supplementary Information [39, 9]. Although other established pipelines [40, 41] exist that offer similar functionality, the `XomicsToModel` pipeline provides several novel capabilities, especially its ability to implement additional constraints, integrate additional data types and relax an infeasible model to recover from the application of inconsistent constraints. These features are primarily enabled by the `thermoKernel` model extraction algorithm.

The mathematical similarities and differences between established model extraction algorithms have been extensively reviewed [42]. Furthermore, a systematic comparative evaluation of model extraction methods [6] has shown that using various data types, such as omics data, for model construction and validation, as well as careful selection of gene expression cut-offs, lead to higher model accuracy when compared with experimental metabolomic data. Subsequently, guidelines for extracting biologically relevant context-specific metabolic models using gene expression data were proposed [43]. The `XomicsToModel` pipeline is implemented in a manner consistent with those guidelines, but the `thermoKernel` model extraction algorithm also adds additional functionality. Table 1 compares `thermoKernel` with other algorithms for extraction of a context-specific model. By default, the `XomicsToModel` pipeline uses `thermoKernel`, but it has been implemented to interface to any of the model extraction algorithms in Table 1 via the generic interface provided by `createTissueSpecificModel` in the COBRA Toolbox [38]. Currently, the `XomicsToModel` pipeline has also been extensively tested with `FastCore` [12] but could readily be interfaced with any other model extraction algorithm.

### Experimental Design

#### Required input data

To run the `XomicsToModel` pipeline, users must first obtain and prepare a generic reconstruction or model and the context-specific data used to extract the model. Furthermore, one must either accept the default set of technical parameters or specify the parameters to suit a particular context (as detailed below).

##### Generic reconstruction or model

A generic genome-scale reconstruction or model represents a metabolic network constructed from the amalgamation and manual literature curation of metabolic reactions that occur in various cell types or organisms. For example, a comprehensive generic reconstruction of human metabolism, such as Recon3D [25], which serves as a valuable resource for metabolic modelling, can be easily accessed and downloaded from the Virtual Metabolic Human database.

##### Context-specific data

Context-specific data represents the genotype or phenotype of a specific biological system. It can be obtained through a review of the existing literature or derived experimentally from a biological system or both. The information can be entered manually or using the function `preprocessingOmicsModel` described in the Supplementary Information. The following context-specific information is currently supported by the `XomicsToModel` pipeline:

- **Bibliomic data:** Data derived from a manual reconstruction following a review of the existing literature. This includes data on the genes, reactions, or metabolites known to be present or absent in the studied biological system. Furthermore, bibliomics data can define a set of coupled reactions or a set of constraints for model reactions based on phenotypic observations.
- **Transcriptomic data:** Measured gene expression levels in the studied biological system. It is used to estimate qualitatively and quantitatively the activity of reactions associated with the detected genes based on the gene-protein-reaction association (GPR; Figure 3). Transcriptomic data can be provided in fragments per kilobase million (FPKM) or raw counts.
- **Proteomic data:** Measured protein levels in the studied biological system. Similarly to transcriptomics data, it can be used to quantitatively estimate which reactions in the metabolic model should be considered active based on the gene-protein-reaction association (Figure 3).
- **Metabolomic data:** The average and standard deviation of metabolite concentrations measured in cell media, biofluids, tissues, or organisms translated into flux units, which may be *mmol*/*gDW*/*h, μmol*/*gDW*/*h* or *nmol*/*gDW*/*h* depending on personal preference. Metabolites detected experimentally can be assigned to be present in the metabolic model. Measured uptake and secretion rates in growth media or biofluids can be used to quantitatively constrain the uptakes and secretions of the model. Furthermore, growth condition information such as growth media composition can also be provided to constrain available uptakes in the model.

Although `XomicsToModel` supports input of all of these different types of context-specific information simultaneously, you only need a subset of these different types of context-specific information to generate a specific model. For example, if there is no proteomic data, it is still possible to generate a context-specific model.

The model includes a set of *technical parameters* that allow users to control the extraction process, adjust boundary conditions, and define how biological and omics data are integrated. These parameters can be modified depending on the analytical objective, but all have default values that ensure the model runs reproducibly under standard conditions. Overall, they can be grouped into five main categories:

- **Boundary and flux control**. These parameters define the allowable flux range and numerical precision for all reactions in the model. Together, these parameters ensure numerical stability and prevent unrealistic reaction rates during optimization.
  - TolMaxBoundary and TolMinBoundary establish the maximum and minimum flux values (default: +1000 and ™1000).
  - fluxEpsilon and thermoFluxEpsilon determine the minimum non-zero flux tolerance (default: primal feasibility tolerance ×10).
  - boundPrecisionLimit refines very small flux values to avoid artificial zeros.
- **Exchange and environmental constraints**. These parameters define how the model interacts with its simulated environment. These options are particularly relevant for reproducing experimental growth conditions or simulating nutrient-limited environments.
  - closeIons and closeUptakes determine whether ion or nutrient uptake reactions are closed (default: *false*).
  - uptakeSign defines the sign convention for uptake fluxes (default: −1).
  - nonCoreSinksDemands (default: ‘closeNone’) specifies whether auxiliary sink or demand reactions remain open.
- **Data integration and biological context**. These parameters control how transcriptomic, proteomic, and metabolomic data are incorporated when generating a context-specific model. These parameters are the most influential for users who wish to adapt the model to a specific dataset or biological condition.
  - transcriptomicThreshold and thresholdP set the minimum expression or abundance levels (logarithmic scale, default: 0).
  - activeGenesApproach defines whether one reaction per active gene (‘oneRxnPerActiveGene’, default) or all corresponding reactions (‘allRxnPerActiveGene’) are included.
  - weightsFromOmics, curationOverOmics, and activeOverInactive determine how literature-curated data and omics-derived weights are prioritized (default: *false*).
  - metabolomicsBeforeExtraction (default: *true*) and metabolomicWeights (default: ‘SD’) control when and how metabolomic data are incorporated.
- **Algorithmic and solver options**. These settings influence the algorithms used for model extraction and optimization. In most analyses, these parameters can remain at their default values unless a specific solver behavior is required.
  - modelExtractionAlgorithm specifies the reconstruction strategy (‘thermoKernel’ by default, ‘fastCore’ as an alternative).
  - fluxCCmethod selects the flux-consistency checking algorithm (‘fastcc’ by default).
  - The relaxOptions structure allows for controlled relaxation in case of infeasibility, while setObjective defines the objective function for optimization.
- **Debugging and output control**. These options are intended for users who need to audit intermediate results or troubleshoot the reconstruction process. For traceability and reproducibility, the model includes:
  - printLevel (verbosity level, default: 0),
  - debug (saves intermediate steps, default: *false*; enabling this can increase storage use),
  - diaryFilename (path to store printed logs).

**Figure 3:**
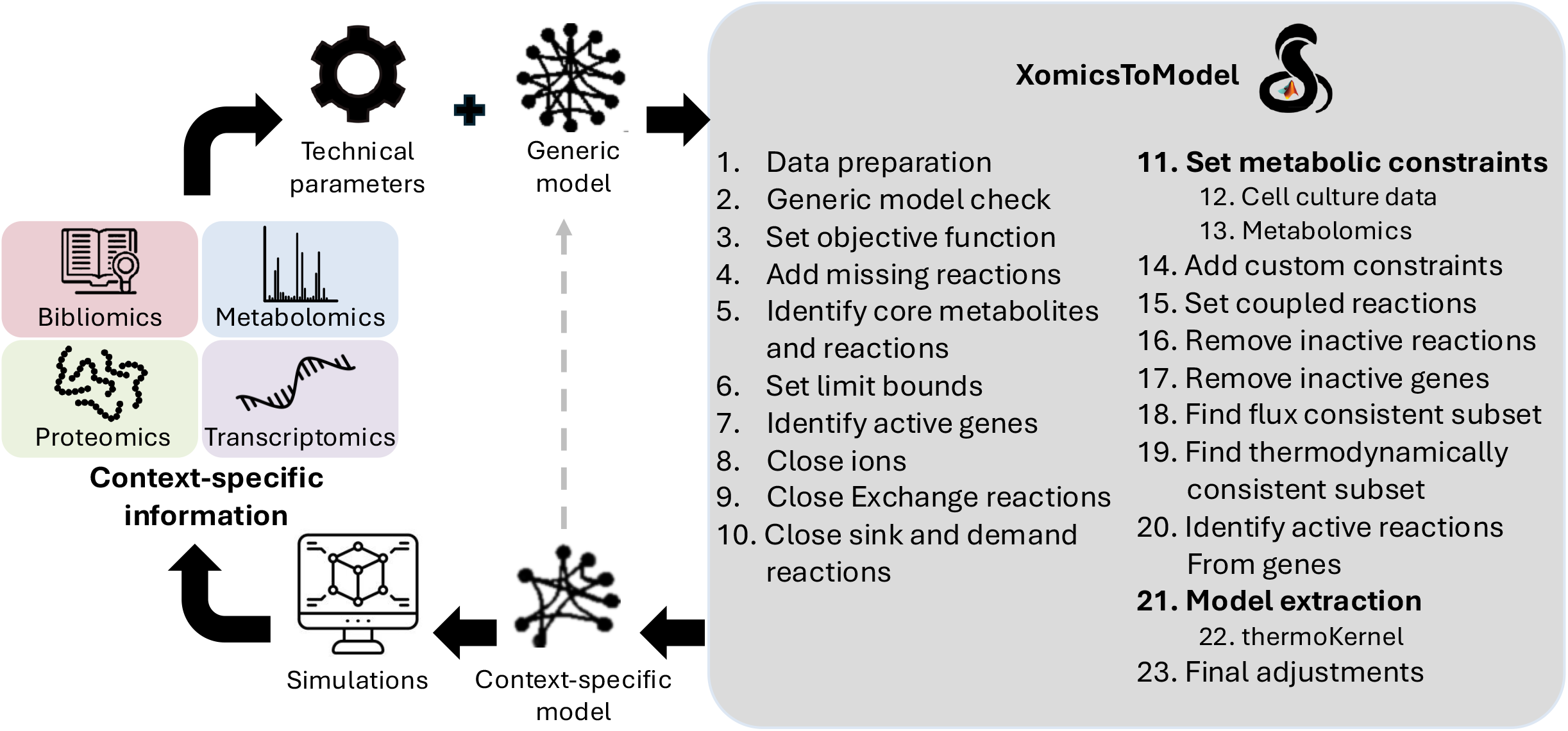
The gene-protein-reaction association or GPRs are boolean operators that describe the interactions of genes, transcripts, proteins, and reactions. For the reaction glucose-6-phosphate isomerase (PGI) to occur, the gene PGI must be activated (Entrez identifier: 2821) leading to the transcription of the PGI mRNA (NCBI Reference Sequence: NM 000175.5). The mRNA is then translated into the glycolytic enzyme PGI (UniProt identifier: P06744). Finally, the PGI enzyme (Enzyme Commission Number: 5.3.1.9) catalyse a reaction that interconverts glucose-6-phosphate (VMH identifier: g6p) and fructose-6-phosphate (VMH identifier: f6p).

##### User guidance

In practice, the parameters most relevant for general users to adjust are:

- Data thresholds and weighting options, which define the biological context of the model.
- Exchange and boundary constraints, which simulate experimental or environmental conditions.

All other parameters can remain at their default values, as they are optimized to ensure numerical robustness, reproducibility, and consistent behavior across analyses.

#### Thermodynamically consistent fluxes (Step 19)

Fluxes must not only satisfy stoichiometric and steady-state constraints but also adhere to the laws of thermodynamics. In this context, a flux is considered thermodynamically feasible if its direction is consistent with a negative Gibbs free energy change, i.e., flux proceeds from higher to lower chemical potential.

A reaction is defined as thermodynamically flux consistent if it admits at least one non-zero flux vector that satisfies both stoichiometric mass balance and thermodynamic feasibility.

Conversely, reactions that can only carry zero flux under all feasible thermodynamic states are classified as thermodynamically inconsistent.

To approximate the largest thermodynamically consistent subset of reactions, `XomicsToModel` employs an iterative strategy that accumulates reactions supporting feasible flux vectors until no further consistent reactions can be identified. This pragmatic approach balances computational tractability with biochemical fidelity.

In practice, Step 19 ensures that only reactions capable of carrying non-zero, thermodynamically feasible fluxes are retained for subsequent model extraction.

Any stoichiometric matrix *S* may be split into one subset of columns corresponding to internal and external reactions, *S* = [*N, B*], where internal reactions are stoichiometrically consistent, that is 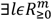 such that *N*^*T*^*l* = 0, and external reactions are not stoichiometrically consistent, that is 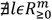 such that *B*^*T*^*l* = 0, as they represent net exchange of mass across the boundary of the system. A steady state flux vector, *v* ∈ *R*^*r*^ such that *Sv* = 0, may be split into an internal reaction flux vector *z* ∈ *R*^*n*^ and an external reaction flux vector *w* ∈ *R*^*k*^, where *Nz* + *Bw* = 0.

We define a thermodynamically feasible flux as a net flux vector where each non-zero entry has a sign opposite change in chemical potential for the corresponding reaction, that is:

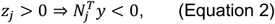

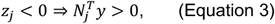

where one may interpret 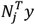 as the thermodynamic driving force for reaction *j*, and interpret *y* ∈ *R*^*m*^ as a vector proportional to the chemical potentials of each metabolite [13]. Since there are fewer metabolites than reactions, any vector of chemical potentials imposes a set of sign constraints on net flues in order for the implications (equation 2) and (equation 3) hold. These equations are a relaxation of the strict thermodynamically feasible flux constraint 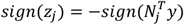, since 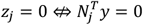.

##### Thermodynamically consistent model

We define a model to be *thermodynamically consistent* if each of its reactions admits a *non-zero* thermodynamically feasible flux. The largest set of reactions that are simultaneously thermodynamically feasible is defined by the cardinality optimisation problem:

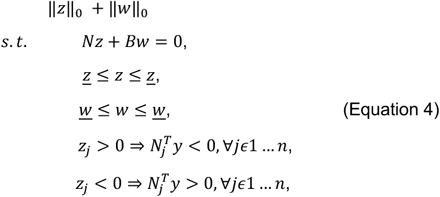

where *z* ∈ *R*^*n*^ is an internal reaction flux vector, *w* ∈ *R*^*k*^ is an external reaction flux vector and *y* ∈ *R*^*m*^ may be interpreted as a vector proportional to the chemical potentials of each metabolite [13]. Here *z* and *z* denote lower and upper bounds on internal reaction fluxes, while *w* ∈ *R*^*k*^ and *w* ∈ *R*^*k*^ denote lower and upper bounds on external reaction fluxes, respectively.

An algorithm for finding the largest thermodynamically feasible subset of a given model and its application to human metabolism is described in detail elsewhere [11]. For completeness, here we reiterate some key points that are relevant to interpretation of this step. Inspired by Desouki et al. [21], we observed that when the bounds on internal reactions constrain the directions but not magnitudes of net flux, that is *z* ∈ {0, −∞}^*n*^ and *z* ∈ {0, ∞}^*n*^, a thermodynamically feasible flux can be computed by a single linear optimisation problem that minimises the absolute value of the internal net reaction flux.

As with flux consistency, typically all reactions in a network do not simultaneously admit a non-zero thermodynamically feasible flux so an approximation to the largest thermodynamically flux consistent subset of reactions in a model can be found by amalgamating reactions with at least one non-zero value in a set of thermodynamically feasible flux vectors. This set is obtained by a sequence of iterations that accumulates thermodynamically flux consistent reactions by biasing net flux toward reactions that have not yet been identified as being thermodynamically flux consistent. This sequence of iterates is terminated when no further thermodynamically flux consistent reactions can be identified or a limit on the number of iterates is reached (by default 30). The constraints in Equation 4 define a non-convex set, so efficiently obtaining an approximation to the largest set of thermodynamically flux consistent reactions is a pragmatic compromise. A convex set is a subset of a vector space (such as Euclidean space) that has a specific property: for any two points within the set, the line segment connecting these points lies entirely within the set. This convexity is crucial for computational efficiency and optimisation in metabolic modelling, providing a robust foundation for refining models and achieving reliable solutions [10].

The last two constraints in Equation 4 are a relaxation of the strict thermodynamic constraint 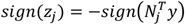since 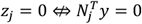. However, they are sufficient to identify a thermodynamically flux consistent subnetwork because any internal reaction with zero flux in all thermodynamically feasible flux vectors, that are each a solution to Equation 4, is defined to be thermodynamically inconsistent.

#### The thermoKernel algorithm (Step 22)

**We designed the** `thermoKernel` **algorithm** to extract a context-specific model that is not only stoichiometrically and flux consistent but also thermodynamically flux consistent.

The core principle of `thermoKernel` is that it balances user-specified evidence of metabolite presence/absence and reaction activity/inactivity with thermodynamic feasibility constraints. The algorithm iteratively identifies feasible flux vectors that maximize inclusion of desired features (e.g., metabolites or reactions) while penalizing inclusion of undesired ones.

Key features of `thermoKernel`:

- Generates models consistent with omics and bibliomic data, weighted by user-specified importance.
- Ensures that all included reactions admit at least one thermodynamically feasible flux.
- Accommodates both qualitative (present/absent) and quantitative (weighted) data.
- Iteratively refines the model to guarantee that every metabolite is produced and consumed, and every retained reaction is active under at least one feasible flux vector.

This algorithm represents a major advance over existing extraction methods by guaranteeing thermodynamic consistency of the resulting models, thereby enabling more reliable predictions of cellular metabolic states.

##### Production and consumption of metabolites

`thermoKernel` assumes that a metabolite is present if it is either produced or consumed at a non-zero rate by a corresponding *net* flux vector, and absent otherwise. Any stoichiometric matrix may be split into *N* = *R* − *F* where *F*_*i*,j_ and *R*_*i*,j_ are the stoichiometric numbers of the *i*^*th*^ molecule consumed and produced in the *j*^*th*^ directed reaction, respectively. In terms of forward reaction flux 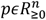 and reverse reaction flux 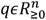, the rate of consumption of each metabolite is *Fp* + *Rq* and the rate of production of each metabolite is *Fq* + *Rp*. Therefore, we observe that *s* ≔ (*F* + *R*)*p* + (*F* + *R*)*q* ≥ 0 represents the sum of the rate of production and consumption of each metabolite. If metabolite i is not present in a network, then *s*_*i*_ = 0.

##### Weighting of metabolites and reactions

With `thermoKernel,` you can generate a context-specific model by providing lists of present metabolites, absent metabolites, active reactions, inactive reactions, and weighted combinations thereof. You can generate a context-specific model only by providing a generic model and a list of present metabolites, a list of active reactions, or a combination of both. With only a list of present metabolites, providing the generic model is thermodynamically flux consistent, a context-specific model is generated containing each of the metabolites required to be present. In this context, there is no constraint on which reactions must be active, except that at least one reaction must be active that results in a non-zero rate of production plus consumption of each metabolite required to be present. With only a list of active reactions, providing the generic model is thermodynamically flux-consistent, a context-specific model is generated containing each of the reactions required to be active. In this context, the list of metabolites that must be present is fully constrained by the list of reactions required to be active. When a combination of lists of present and absent metabolites as well as active and inactive reactions is provided, the resulting context-specific model is a trade-off between these specifications, where by default equal weight is given to metabolite presence, metabolite absence, reaction activity and reaction inactivity.

##### Model extraction

`thermoKernel` identify a context-specific yet thermodynamically consistent model from a model that is flux, or also thermodynamically flux-consistent. Each iteration of `thermoKernel` identifies a thermodynamically feasible flux vector that simultaneously quantitatively balances incentives for the presence of metabolites and the activity of reactions, with disincentives for absence of metabolites and inactivity of reactions, utilising the following optimisation approach. This is represented by the following optimisation problem:

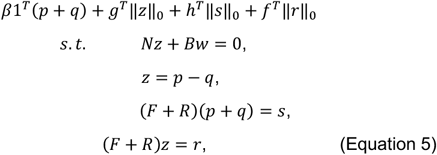

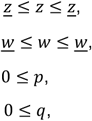

Equation 5 contains the following variables:

- *zϵR*^*n*^ is an internal reaction flux vector.
- *wϵR*^*k*^ is an external reaction flux vector.
- 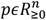is forward net reaction flux.
- 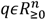is reverse net reaction flux.
- 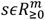m is the sum of the rate of production and consumption of each metabolite.
- *rϵR*^*m*^ is an approximation to the sum of production and consumption of each metabolite due to net reaction flux.

Equation 5 contains the following data matrices:

- *NϵN*^*m*×*n*^ and *BϵN*^*m*×*k*^ are internal (stoichiometrically consistent) and external (stoichiometrically inconsistent) stoichiometric matrices.
- *FϵN*^*m*×*n*^ and *RϵN*^*m*×*n*^ are forward and reverse stoichiometric matrices [44], with *F* = *max*(−*N*, 0).
- *R* = *max*(*N*, 0) and *N* = *R* − *F*.

Equation 5 contains the following data vectors

- *z* ∈ {0, −∞}^*n*^ and *z* ∈ {0, ∞}^*n*^ denote lower and upper bounds on internal reaction fluxes respectively, while
- *wϵR*^*k*^ and *w∈R*^*k*^ denote lower and upper bounds on external reaction fluxes, respectively.

In Equation 5, the objective term *β*1^*T*^(*p* + *q*) minimises the absolute value of net internal reaction flux *zϵR*^*n*^, which optimises toward a thermodynamically feasible flux vector [21]. The scalar weight *β* > 0 is chosen to trade-off between cardinality optimisation of internal reaction fluxes, which explores the flux consistent subset of the model, and minimising the one norm of the sum of forward and reverse reaction rates, which promotes thermodynamic feasibility. The parameter vector *gϵR*^*n*^ is a vector of weights on internal reaction rates. The term *g*^*T*^‖*z*‖_0_ denotes the weighted zero-norm of internal reaction fluxes, where the zero-norm for each reaction is weighted individually, that is:

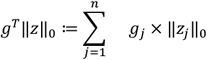

When *g*_*j*_ < 0, or *g*_*j*_ > 0, then net flux is incentivised or penalised, in reaction *j*, respectively. In Equation 5, the parameter vectors *h, f∈R*^*m*^ are vectors of weights on metabolites. When *h*_*j*_ > 0, the ob*j*ective term 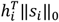minimises the zero-norm of the rate of production plus consumption of metabolite *i*, promoting its elimination from a model. When *h*_*j*_ < 0, the objective term 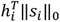maximises the zero-norm of the rate of production and consumption of metabolite *i*. However, even if *s*_*i*_ > 0, then a metabolite may not be produced or consumed at a non-zero rate by a corresponding *net* flux vector since *p* = *q* ≠ 0 is a solution that also implies *s* > 0. Therefore, we introduce the variable *r* ≔ (*F* + *R*)*z* as an approximation to the sum of production plus consumption due to net reaction flux, which is (*F* + *R*)|*z*|, but is non-trivial to implement. If *r*_*i*_ ≔ (*F* + *R*)_*i*_*z* > 0 then (*F* + *R*)_*i*_|*z*| > 0 as desired. Therefore, when *f*_*i*_ < 0, the ob*j*ective term 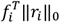approximates maximisation of the weighted zero-norm of the sum of production plus consumption of metabolite *i* due to non-zero net reaction flux. In practice, we set *f*_*i*_ = −*min* (*h*_*i*_, 0) so that optimisation of the cardinality of *s*_*i*_ and *r*_*i*_ is concordant.

In Equation 5, the internal bounds *z* ∈ {0, −∞}^*n*^ and *z* ∈ {0, ∞}^*n*^ enforce reaction directionality without interfering with minimisation of the absolute value of internal net flux. That is, if *g* = *h* = *f* = 0 then Equation 5 is guaranteed to return a thermodynamically feasible flux. However, we find that with *g* = *h* = *f* ≠ 0, Equation 5 still returns a thermodynamically feasible flux provided that the value of the strictly positive scalar parameter *β* is such that *β*1^*T*^(*p* + *q*) ≥ *g*^*T*^‖*z*‖_0_ + *h*^*T*^‖*s*‖_0_ + *f*^*T*^‖*r*‖_0_ at the optimum of Equation 5. In this way, Equation 5 optimises toward thermodynamically feasible flux and simultaneously quantitatively trades off incentives for presence of metabolites and activity of desired reactions, with penalties for inclusion of undesired metabolites and reactions.

The following optional variants of Equation 5 are implemented in `thermoKernel` depending on the data included in the preceding steps:

- A linear objective (Step 3) on net flux may be added to the objective in Equation 5 to optimise a linear combination of net fluxes, as in flux balance analysis [14]. This will not interfere with thermodynamic feasibility if all the reactions being optimised are stoichiometrically inconsistent, i.e., exchange reactions. However, it may be that application of thermodynamic feasibility constraints reduces the maximum value of the part of the objective that optimises net flux.
- Coupling constraints [35] (Step 15), *Cv* ≤ *d*, may be added to Equation 5 but depending on how they are formulated, such constraints may interfere with the ob*j*ective term *β*1^*T*^(*p* + *q*) that optimises toward thermodynamically feasible flux. In that case, the coupling constraints may render part of the extracted model thermodynamically infeasible (e.g., coupling constraints may be formulated that force net flux around a stoichiometrically balanced cycle).
- The box constraints net flux in Equation 5 may be used to force non-zero flux in a particular direction of one or more internal reactions, that is *z, z∈R*^*n*^. Depending on the choice of bounds (Step 14), this may result in extraction of a model where some reactions are thermodynamically inconsistent (e.g., a combination of bounds that force net flux around a stoichiometrically balanced cycle).
- In `thermoKernel,` the implementation of Equation 5 and its variants uses the default *β* ≔ 1*e* − 4. By setting `model.beta`, you can set a larger value of *β*, which will increase the incentive to find a fully thermodynamically feasible flux vector in each iteration of Equation 5 and decrease the incentive to search the steady state solution space for a flux vector that results in certain reactions and metabolites to be active and present, respectively. However, if *β* is too large, relative to the magnitudes of the weights on metabolites and reactions, this impedes the ability of Equation 5 to search the full set of thermodynamically feasible fluxes.
- Similarly, there is a trade-off between the reaction weights *gϵR*^*n*^ and the metabolite weights *h, fϵR*^*m*^, where we set *f*_*i*_ = −*min* (*h*_*i*_, 0), so their relative magnitude should reflect the relative importance of the corresponding input bibliomic, transcriptomic and proteomic data (Step 7).

Although it is not always necessary, the default `thermoKernel` implementation for each iteration of Equation 5 should be followed by the linear optimisation problem first introduced in previous work [21]. This minimises the absolute value of internal reaction flux, while (a) all exchange fluxes are kept constant, and (b) no internal flux is allowed to change direction or increase in size. This has two advantages. Firstly, it is an efficient test to check that Equation 5 returns a solution where every reaction corresponds to a thermodynamically feasible flux. Secondly, it can be used to remove that part of a flux vector that corresponds to flux around a stoichiometrically balanced cycle and the output is a flux vector that is thermodynamically feasible, i.e., each reaction flux satisfies equations 2 and 3. When `XomicsToModel` calls `thermoKernel`, the default behaviour is for flux bounds, which interfere with thermodynamic feasibility, to be relaxed when testing for cyclic flux after Equation 5. This prioritises thermodynamic consistency of models generated by `XomicsToModel` over user input constraints that attempt to force a contradiction.

##### Extracting a context-specific and thermodynamically consistent model

Given a generic model, it is usually the case that more than one thermodynamically feasible flux vector is required to ensure the activity of a specified subset of its reactions, presence of a specified subset of its metabolites, or both. That is, a single thermodynamically feasible flux vector is usually unable to enforce the activity and presence of all the reactions and metabolites in a model. Therefore, a randomly greedy strategy is implemented, inspired by previous work [45], whereby it chooses a random vector from a uniform rectangular set, *d* ∈ {*R*^*n*^|*unif*(0,1)}, then updates the weights on cardinality optimisation of reactions in the next iterate *g*(*n* + 1) based on those of the previous iterate *g*(*n*), with the heuristic:

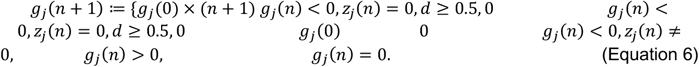

This approach successively increases the incentive for activity of internal reactions that are desired to be active but have not yet been active in a previous iterate. The strategy is the same to incentivise presence of metabolites. The iterations either conclude when all incentivised reactions and metabolites are active and present, respectively, or when a prespecified maximum number of iterations is reached. Since reaction *j* is defined to be active when *z*_*j*_ ≠ 0 (numerically implemented by requiring that |*z*_*j*_| ≥ *ε* with *ε* = 1*e* − 4 in practice), every reaction added to a thermodynamically consistent model is a reaction that admits at least one non-zero thermodynamically feasible flux and every metabolite is produced and consumed by at least one thermodynamically feasible flux. Therefore, while generating a single thermodynamically feasible flux vector that containing non-zero and zero fluxes, the strict thermodynamically feasible flux constraint 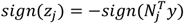is relaxed; a thermodynamically consistent model is only generated from reactions that admit non-zero thermodynamically feasible fluxes. Therefore, the model generated is fully consistent with thermodynamics in its broadest sense. Each iteration of Equation 5 is efficiently solved using a piecewise linear sequence of linear optimisation problems, as described in detail elsewhere [11].

### Limitations

This protocol focuses on extracting a context-specific genome-scale model from an existing reconstruction by integrating omics data from a biological system. It does not cover generation of a generic, genome-scale metabolic reconstruction from scratch [4, 2], or the analysis of existing models [38]. Additionally, the integration of omics data presents several challenges since they are generated using a variety of platforms, affecting storage and data formats significantly. The integration of omics data necessitates data in specific formats. Therefore, individual omics data must be pre-processed. Furthermore, experimental errors, such as data processing or measurement errors, can propagate through the extraction of a metabolic network that is not a faithful representation of the original biochemical system. In this situation, a significant loss of predictive capacity will be evident. Therefore, it is not recommended to extract a context-specific model and claim that it has high predictive accuracy without commensurate comparison with some independent experimental data.

We define a steady-state flux vector to be thermodynamically feasible if it is consistent with energy conservation and the second law of thermodynamics [16, 17]. They are thermodynamic constraints that apply to chemical and biochemical systems alike. In addition, biochemical systems are subject to many additional constraints that range from abiotic constraints [46] to constraints imposed by evolutionary pressures toward optimization [47]. For example, there are biochemical thermodynamic and enzyme kinetic constraints on the range of metabolite concentrations required for life [25]. Limits to the feasible range of metabolite concentrations lead to additional constraints on the direction [48] and magnitude [49]of reaction flux that are not applied within the `XomicsToModel` pipeline. The models generated by `XomicsToModel` are thermodynamically flux consistent, they can be integrated with additional constraints arising from biochemical thermodynamics and kinetics without a risk of generating a thermodynamically infeasible model. Several studies support this approach. For instance, one study [50] demonstrated the necessity of thermodynamic consistency in cyclic enzyme-catalysed reaction networks, highlighting the strong constraints on kinetic parameters imposed by zero net flux cycles. Other groups [51] developed a formalism to combine kinetic and thermodynamic measurements into state-resolved models that are thermodynamically consistent, correcting potential inconsistencies in the models. These approaches ensure that the models are both accurate and feasible within the constraints of thermodynamic laws, supporting the assertion that `XomicsToModel` can handle additional biochemical constraints without issues.

## MATERIALS

### Equipment

#### Input data

The COBRA Toolbox supports commonly used data formats for model description such as Systems Biology Markup Language (SBML) and Excel sheets (.xlsx). MATLAB supports a wide range of text and spreadsheet formats that can be used to provide context-specific data for the `XomicsToModel` pipeline.

#### Required hardware

- A computer with at least 8 GB of RAM and any 64-bit Intel or AMD processor. **▲ CRITICAL** Depending on the size of the model, more processing power and more memory might be needed.
- A hard drive with at least 10 GB of space available.

#### Required software

- An operating system that is MATLAB qualified (https://mathworks.com/support/sysreq.html). **▴ CRITICAL** To make sure that the operating system is compatible with the MATLAB version, check the requirements at https://mathworks.com/support/sysreq/previous_releases.html.
- MATLAB (MathWorks, https://mathworks.com/products/matlab.html), version R2021+. Install MATLAB and its license by following the official installation instructions (https://mathworks.com/help/install/ug/installmathworks-software.html). **▴ CRITICAL** Version R2021+ or above is recommended for running MATLAB live scripts (.mlx files). **▲ CRITICAL** No support is provided for versions older than R2021. MATLAB is released on a twice-yearly schedule, and the latest releases of MATLAB may not be compatible with the existing solver interfaces, necessitating an update of the MATLAB interface provided by the solver developers, an update of the COBRA Toolbox, or both.
- The COBRA Toolbox version 3.4 [38], or above. To install the COBRA Toolbox, follow the instructions on https://github.com/opencobra/cobratoolbox. **▴ CRITICAL** If an installation of COBRA Toolbox is already present, update the repository from MATLAB, via terminal, or Git Bash. **▲ CRITICAL** Check that all of the system requirements in https://opencobra.github.io/cobratoolbox/docs/requirements.html are met.

#### Solvers

The table below provides an overview of the optimisation solvers supported by the `XomicsToModel` pipeline.

| Name | Version | Interface |
| --- | --- | --- |
| GLPK | 2.7+ | glpk |
| GUROBI | 7.0+ | gurobi |
| ILOG CPLEX | 10-12.10 | ibm_cplex |
| MATLAB | R2014b+ | matlab |
| MOSEK | 8.0+ | mosek |

### Equipment setup

#### COBRA Toolbox

Unless you are already a developer of COBRA Toolbox code, update the COBRA Toolbox from within MATLAB by running the following command:

`>> updateCobraToolbox`

Alternatively, update from the terminal (or Git Bash) by running the following from within the COBRA Toolbox directory.

`$ cd cobratoolbox # change to the local cobratoolbox directory`

`$ git checkout master # switch to the master branch`

`$ git pull origin master # retrieve changes`

If the update of the COBRA Toolbox fails or cannot be completed, clone the repository again.

**▲ CRITICAL** The COBRA Toolbox can be updated as described above only if no changes have been made to the code in one’s local clone of the official online repository. To contribute any local edits to the COBRA Toolbox code into the official online repository, please follow the MATLAB.devTools guidelines [38] (https://opencobra.github.io/MATLAB.devTools/stable/).

To generate a context-specific model from a generic model (or reconstruction) using omics data, first launch the COBRA Toolbox [38] and specify the optimisation solver to be used.

`>> initCobraToolbox`

`>> changeCobraSolver(‘gurobi’, ‘LP’);`

`>> changeCobraSolver(‘gurobi’, ‘QP’);`

## PROCEDURE

**▴ CRITICAL** The sequence of steps described below are in the same order as they are implemented by `XomicsToModel`. This is important because the overall approach to model generation is to first increase the dimensions and magnitude of the feasible set of steady-state metabolic fluxes, then successively decrease its dimensions and magnitude, until a context-specific model and information about the procedure and result of the model generation process is returned to the user after the final step. The `XomicsToModel` pipeline can be run if at least one input variable and a generic model (in standard COBRA Toolbox format https://github.com/opencobra/cobratoolbox/blob/master/docs/source/notes/COBRAModelFields.md) is provided. The user-defined context-specific information (`specificData`) and technical parameters (param) are optional, and if not specified, default values for the required fields will be assigned.

`>> [contextSpecificModel, modelGenerationReport] = XomicsToModel(genericModel, specificData, param)`

The name and types of the fields for the `specificData` and param variables must be identical to how they are described in each step of the `XomicsToModel` pipeline below, so the `XomicsToModel` pipeline can recognise them. The supplementary `preprocessingOmicsModel` function can be used to generate the `specificData` variable, as described in the Supplementary Information.

### Data Preparation

**CRITICAL** In this first step, `XomicsToModel` checks the context-specific data in `specificData`, checks the technical parameters in param, and checks the generic model to test if they are compatible with the naming conventions and formats expected by the rest of the `XomicsToModel` pipeline. `XomicsToModel` identifies missing `specificData` and param fields, and default values are assigned for optional inputs.

**1**| To aid interpretation of the performance of the pipeline, specify a filename for the diary in `param.diaryFilename`. To enhance the interpretation of the `XomicsToModel` pipeline performance, a verbose level can be implemented in `param.printLevel`.

To control the amount of information displayed for each step, adjust the verbose level parameter from 0 to 3. A verbose level of 0 provides minimal output, focusing only on essential information (but not completely silent), while a level of 1 includes additional warnings and status updates. At level 2, more detailed process information is displayed, including intermediate steps, and at level 3, the most comprehensive output is provided, showing detailed logs and debugging information.

Within this diary file, many intermediate results of the pipeline are printed that are essential for debugging. If a certain reaction is missing that was expected within the generated model, search for it in the diary file. Specify `param.debug = 1` to print out intermediate results after key steps of the `XomicsToModel` pipeline. Later steps in the Procedure will mention that results are printed; to find these results, open the diary.

These intermediate results can be re-loaded so that subsequent individual steps of the `XomicsToModel` pipeline may be run to debug unexpected results. Alternatively, these intermediate results can be analysed as a batch to monitor the model generation progress, as described in the Supplementary Information.

#### Usable Variables for Step 1

`param.debug:` Logical, true if the pipeline should save its progress for debugging (Default: false).

`param.diaryFilename:` The name (and location) of a diary file with the printed pipeline output (Default: 0).

`param.printLevel:` Level of verbose that should be printed (Default: 0).

### Generic model checks

**CRITICAL** In this step, the generic model is checked for compatibility with `XomicsToModel`. It is checked for fields that may cause inconsistency when reactions are added or removed by `XomicsToModel`. These checks depend on the model extraction algorithm used by `XomicsToModel`.

**2**| Select your preferred model extraction algorithm by setting `param.tissueSpecificSolver`. The default is `‘thermoKernel’`, which will ensure the model extracted admits a thermodynamically feasible flux for each reaction. Alternatively, set param.tissueSpecificSolver = ‘fastcore’.

Regardless of the model extraction algorithm selected, every generic model must satisfy the steady state constraints *Sv* = 0, reaction rate bounds *l* ≤ *v* ≤ *u*, and optionally coupling constraints *Cv* ≤ *d*, implying that the system of inequalities is feasible; if the model is infeasible in this step, an error will be generated Review the log; if `param.printLevel` is greater than zero, the information in `specificData` and `param` will be printed.

#### ? TROUBLESHOOTING

##### Usable Variables for Step 2

`param.printLevel:` Level of verbose that should be printed (Default: 0). `param.tissueSpecificSolver:` The name of the tissue-specific solver to be used to extract the context-specific model (Possible options: `‘thermoKernel’` and `‘fastcore’`; Default: `‘thermoKernel’`).

### Set objective function (optional)

**CRITICAL** `XomicsToModel` has the capability to set an objective function as used in Flux Balance Analysis [14], where *c* ∈ *R*^*n*^ represents the biologically inspired linear ob*j*ective to find the optimal flux vector.

**3**| (Optional) Unless you are using default settings, set the linear objective *φ*(*v*) ≔ *c*^*T*^*v* in `param.setObjective.` The objective function in param.setObjective represents the reaction whose flux is to be maximised (such as ATP or biomass production) or minimised (such as energy consumption). However, alternative biological objectives can also be used. For example, these may include coupled reactions or mathematical objectives like the minimisation of zero, one, or two norms, entropy maximisation, or a combination of biological and mathematical objectives. Examples of such combinations include the minimisation of norms combined with reaction expression data based on gene expression or the minimisation of bonds broken and formed based on atom mapping data at a genome scale. These approaches are illustrated in Preciat et al. [9].

#### Usable Variable for Step 3

`param.setObjective:` Reaction identifier indicating the reaction to maximise (Default: none).

### Add missing reactions (optional)

**▲ CRITICAL** If a reaction is already present in the draft model, it will be replaced by the new reaction list.

**▲ CRITICAL** By associating a new gene identifier with a reaction, a new gene will be added to the draft model.

**4**| (Optional) If reactions are missing from the generic model, specify the missing reactions to add in `specificData.rxms2add`. These reactions are then automatically added to the draft model by `XomicsToModel` using the function [addReaction](https://opencobra.github.io/cobratoolbox/stable/modules/reconstruction/refinement/index.html?highlight=addreaction#src.reconstruction.refinement.addReaction). When specifying missing reactions, multiple data can be included but only the reaction identifier and the reaction formula are mandatory.

Add missing metabolic reactions on `specificData.rxns2add`. The default bounds are set based on the arrows used in the reaction formula to separate the substrates and products as shown in the table `specificData.rxns2add` in ‘Usable Variables for Step 4’ (below). The available arrows are:

- Forward (−>): *l*_*j*_ = 0; *u*_*j*_ = `param.TolMaxBoundary`.
- Reverse (< −): *l*_*j*_ = `param.TolMinBoundary`; *u*_*j*_ = 0.
- Reversible (<=>) *l*_*j*_ = `param.TolMinBoundary`; *u*_*j*_ = `param.TolMaxBoundary`.

If not otherwise specified, the default values for missing reaction names, metabolic pathways, and gene rules are *Custom reaction, Miscellaneous*, and an empty cell, respectively.

Adding reactions to a model may render it stoichiometrically inconsistent. Given a generic reconstruction or model, `XomicsToModel` generates a generic stoichiometrically consistent model using the function [findStoichConsistentSubset](https://opencobra.github.io/cobratoolbox/stable/modules/reconstruction/modelGeneration/stoichConsistency/index.html?highlight=findstoichconsistentsubset#src.reconstruction.modelGeneration.stoichConsistency.findStoichConsistentSubset) described elsewhere [11]. Briefly, this function approximately solves the problem to extract the largest subset of a reconstruction or model that is stoichiometrically consistent [38]. A set of reactions is stoichiometrically consistent if every reaction in that set is mass balanced [52]. The stoichiometric matrix may be split *S* ≔ [*N, B*], where the matrix *N* represents stoichiometrically consistent internal reactions, while the matrix *B* represents stoichiometrically inconsistent external reactions.

Review the log; if `param.printLevel` is greater than zero, a table with a summary of the draft model’s stoichiometric consistency will be printed (Table 3).

Inconsistent metabolites and reactions are removed; if this renders a model infeasible with respect to steady state flux, an error is generated by `XomicsToModel`. Also in this step, the gene-protein-reaction rules are updated for the newly added reactions, and the reaction-gene-matrix is regenerated. In addition, different boolean vectors indicating stoichiometrically consistent and inconsistent metabolites and reactions are added to the draft model.

**▲ CRITICAL STEP** Extracting the largest stoichiometrically consistent subset is combinatorially complex. At genome-scale, only approximation to the actual solution is achievable in practice. The quality of the approximation is improved if `findStoichConsistentSubset` is warm-started with reactions specified as being stoichiometrically inconsistent (e.g., external reactions specified as false in the corresponding entry in `model.SIntRxnBool)`. If `model.SIntRxnBool` is not provided, `XomicsToModel` calls the function `findSExRxnInd` to heuristically distinguish internal and external reactions based on reaction naming conventions (e.g. ‘EX_’ prefix for an exchange reaction) and reaction stoichiometry (any reaction with only one non-zero stoichiometric coefficient in the corresponding row is an external reaction).

#### ? TROUBLESHOOTING

##### Usable Variables for Step 4

`model.SIntRxnBool:` Logical vector, true if a reaction is heuristically considered an internal reaction, false for an external reactions (Default: false). `specificData.rxns2add:` Table containing the identifier of the reaction to be added, its name, the reaction formula, the metabolic pathway to which it belongs, the gene rules to which the reaction is subject, and the references. (Default: empty).

| rxns | rxnNames | rxnFormulas | subSystems | rxnGrRules | rxnReferences |
| --- | --- | --- | --- | --- | --- |
| newRxn1 | Oxidation | A + O -> AO | Glycolysis | gene1 or gene2 | PMID: **** |
| newRxn2 | Reduction | AH + B <=> A + BH | Glycolysis | gene1 | PMID: **** |

`param.TolMaxBoundary:` The reaction flux upper bound maximum value (Default: 1000).

`param.TolMinBoundary:` The reaction flux lower bound minimum value (Default: −1000).

### Identify core metabolites and reactions (optional)

**CRITICAL** The `XomicsToModel` pipeline uses several user-specified fields from the `specificData` variable to identify the set of metabolites and reactions that must be active in the draft model. These optional fields include the reaction specified as the objective function, the metabolic reactions to add, the metabolites and reactions identified as active based on bibliomic data, the reactions whose bounds are restricted based on bibliomic data, the coupled reactions, the exchange reactions for metabolites present in the growth media, and the exchange reactions for metabolites shown to be secreted or taken up based on exometabolomic data.

**5**| (Optional) Add reactions to the active reaction list, which will be appended or shortened if necessary to keep the model stoichiometrically consistent, flux consistent (and optionally thermodynamically consistent). To configure these fields, perform the following actions and refer to the Usable Variables table for syntax:

- Activate specific reactions in `specificData.activeReactions` based on bibliomic data.
- Specify a linear objective function in `param.setObjective` (refer to Step 3).
- Generate a table of reactions to be added in `specificData.rxns2add` (refer to Step 4).
- List the metabolites known to be absent in `specificData.absentMetabolites` based on the bibliomic data.
- List of metabolites known to be present in `specificData.presentMetabolites` based on bibliomic data.
- Include a list of metabolites present in the cell culture in `specificData.mediaData` if the omics data originated from one.
- Utilize qualitative or semi-quantitative exometabolomic data in `specificData.exomet`
- Formulate a list of reactions for constraint in `specificData.rxns2constrain` based on bibliomic data.
- Establish a set of coupled reactions in `specificData.coupledRxns`

#### Usable Variables for Step 5

`param.setObjective:` Cell string indicating the linear objective function to optimise (Default: none).

`specificData.activeReactions:` List of reactions know to be active based on bibliomic data (Default: empty).

`specificData.coupledRxns:` Table containing information about the coupled reactions. This includes the identifier for the coupling constraint (couplingConstraintID), the list of coupled reactions (coupledRxnsList), the coefficients of those reactions (c, given in the same order), the right hand side of the constraint (d), the constraint sense or the directionality of the constraint (dsence), and the reference (Default: empty).

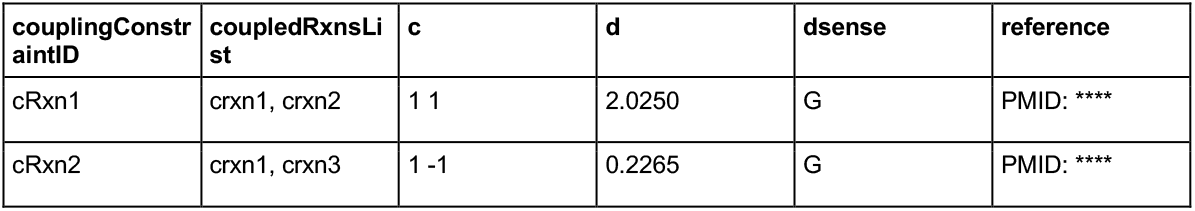

| couplingConstraintID | coupledRxnsList | c | d | dsense | reference |
| --- | --- | --- | --- | --- | --- |
| cRxn1 | crxn1, crxn2 | 1 1 | 2.0250 | G | PMID: **** |
| cRxn2 | crxn1, crxn3 | 1 -1 | 0.2265 | G | PMID: **** |

`specificData.exomet:` Table with measured exchange fluxes, e.g., obtained from an exometabolomic analysis of fresh and spent media. It includes the reaction identifier, the reaction name, the measured mean flux, standard deviation of the measured flux, the flux units, and the platform used to measure it (Default: empty).

| rxns | rxnNames | mean | SD | units | platform |
| --- | --- | --- | --- | --- | --- |
| EX_mMet1 | Exchange of mMet1 | 100 | 8x10<sup>-4</sup> | μ mol/gDW/h | LC-MS |
| EX_mMet2 | Exchange of mMet2 | 4x10<sup>-3</sup> | 3x10<sup>-4</sup> | μ mol/gDW/h | GC-MS |

`specificData.mediaData:` Table containing the fresh media concentrations. Contains the reaction identifier, the maximum uptake (μ mol/gDW/h) assuming it is equal to the concentration of the metabolite in fresh medium divided by the length of time before medium exchange, and the medium concentration (μ mol/l; Default: empty).

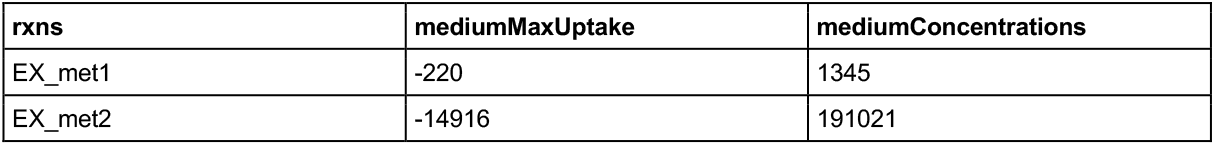

| rxns | mediumMaxUptake | mediumConcentrations |
| --- | --- | --- |
| EX_met1 | -220 | 1345 |
| EX_met2 | -14916 | 191021 |

`specificData.rxns2add:` Table containing the identifier of the reaction to be added, its name, the reaction formula, the metabolic pathway to which it belongs, the gene rules to which the reaction is subject, and the references. (Default: empty).

| rxns | rxnNames | rxnFormulas | subSystems | rxnGrRules | rxnReferences |
| --- | --- | --- | --- | --- | --- |
| newRxn1 | Oxidation | A + O -> AO | Glycolysis | gene1 or gene2 | PMID: **** |
| newRxn2 | Reduction | AH + B <=> A + BH | Glycolysis | gene1 | PMID: **** |

`specificData.rxns2constrain:` Table containing the reaction identifier, the updated lower bound (lb), the updated upper bound (ub), a constraint description, and any notes such as references or special cases (Default: empty).

| rxns | lb | ub | constraintDescription | Notes |
| --- | --- | --- | --- | --- |
| rxn1 | 0 |  | Close uptake | PMID: **** |
| rxn2 | 0.0.956 | 1000 | Turnover constraints | PMID: **** |

### Set limit bounds (optional)

**6**| (Optional) Considering the constraints *l* ≤ *v* ≤ *u* shown in Equation 1, where *l* is the lower bound and *u* the upper bound, set `param.TolMinBoundary` so that any minimum lower bound is assigned to the minimum value of the lower bounds. Similarly, set `param.TolMaxBoundary` so that any maximum upper bound is assigned to the maximum value of the upper bounds. After this adjustment, (*l*) = *param. TolMinBoundary* and (*u*) = *param. TolMaxBoundary*.

To modify the lower and upper boundaries of the reaction fluxes, set:

- `param.TolMinBoundary =` *min*(*l*).
- `param.TolMaxBoundary =` *max*(*u*).

`param.TolMinBoundary` represents the minimum value a reaction can have, ensuring that no reaction in the network will have a flux lower than this threshold. Similarly, `param.TolMaxBoundary` sets the maximum allowable flux value for reactions, preventing fluxes exceeding this threshold in the entire network. The model’s feasibility is tested with the new bounds.

Review the log; if `param.printLevel` is greater than zero, the modified bounds will be printed.

#### Usable Variables for Step 6

`param.TolMaxBoundary:` The reaction flux upper bound maximum value (Default: 1000).

`param.TolMinBoundary:` The reaction flux lower bound minimum value (Default: −1000).

**▲ CRITICAL STEP** There should be no lower bound that exceeds an upper bound.

### Identify active genes (optional)

**▲ CRITICAL** A gene must be present in the draft model to be considered active.

**7**| Specify a list of active genes by listing their NCBI Entrez gene identifiers, which are unique integers for genes or other loci, or by using the gene identifier employed in the generic model. Compose the active genes list based on data from multiple sources, such as Bibliomics, Proteomics and/or Transcriptomics:

- Specify which genes should be added using the gene-protein-reaction rules in `specificData.rxns2add.rxnGrRules` (Step 4).
- Bibliomics: Classify genes as active in `specificData.activeGenes` based on curation of published literature using your preferred specifications.
- Proteomics: Classify genes as active based on their (normalised) proteomic peak areas from the list of detected proteins (`specificData.proteomics`). If *peakArea* ≥ `param.thresholdP`, the gene is considered active.
- Transcriptomics: Add on `specificData.transcriptomics` genes that, based on the list of detected transcripts, are active based on their expression levels (FPKM, Fragments Per Kilobase of transcript per Million mapped reads or raw counts). If *log*(*expressionLevel*) ≥ `param.thresholdT`, the gene is considered active.

The parameter `inactiveGenesTranscriptomics` determines whether reactions associated with genes whose transcript levels fall below “`param.thresholdT`”, are removed from the draft model. Setting this parameter to “`true`” excludes these reactions, refining the model to include only transcriptionally supported metabolic functions. Conversely, setting it to `false` retains all reactions regardless of transcript levels, which may be preferred when transcriptomic data are incomplete or when integrating multiple omics datasets.

In the event of a discrepancy, such as an inactive gene according to manual literature curation but active according to proteomic or transcriptomic data:

- Set `param.curationOverOmics = true` to give preference to bibliomic data.
- Set `param.curationOverOmics = false` to give preference to omics data.

By default, bibliomic data takes precedence over other omics data, i.e., `param.curationOverOmics = false`. Genes not found in the draft model are removed from the list of active genes.

Furthermore, two fields are added to the model to indicate the gene expression value of the genes present in the draft model `model.geneExpVal` and the expression of the corresponding reactions (`model.expressionRxns`) based on the function `mapExpressionToReactions`. The logarithm of the levels of gene expression in `specificData.transcriptomicData` are used to link gene expression levels to associated reactions using the function `mapExpressionToReactions`. In short, the resulting `model.expressionRxns` field is based on the gene relationship described in the gene-protein-reaction rules: an **AND** in the gene-protein-reaction rules is assigned *min*(*l*), and an **OR** is assigned a *max*(*u*). For genes that are not found in the transcriptomics data, no expression value is added (NaN is the default).

Although the default threshold for specificData.proteomics and specificData.transcriptomicData data is 0, a common practice in the field is to set it to *log*(2), corresponding to a fold change of 2. This threshold indicates that the expression level of a gene or protein is at least doubled compared to the baseline, highlighting a biologically meaningful increase and reducing noise from low-expressed genes or proteins [53].

Review the log; if `param.printLevel` is greater than zero, a histogram of reaction expression is printed.

Proteomics data is usually annotated with UniProtKB. To integrate proteomics data with a generic model, it is essential to convert UniProtKB identifiers to gene identifiers using bioinformatics tools such as DAVID or UniProt’s mapping services. This conversion allows seamless translation and facilitates comprehensive analyses across diverse biological databases.

#### Usable Variables for Step 7

`param.curationOverOmics:` Logical, indicates whether curated data should take priority over omics data (Default: false).

`param.inactiveGenesTranscriptomics:` Logical, indicate if inactive genes in the transcriptomic analysis should be added to the list of inactive genes (Default: true). `param.thresholdP:` The proteomic cutoff threshold for determining whether or not a gene is active (Default: 0).

`param.thresholdT:` The transcriptomic cutoff threshold for determining whether or not a gene is active (Default: 0).

`specificData.activeGenes:` List of Entrez identifier of genes that are known to be active based on the bibliomic data (Default: empty).

`specificData.inactiveGenes:` List of Entrez identifier of genes known to be inactive based on the bibliomics data (Default: empty).

`specificData.proteomics:` Table with a column with Entrez ID’s and a column for the corresponding protein levels (Default: empty).

| gene | expVal |
| --- | --- |
| entrezID1 | 24 |
| entrezID2 | 78 |

`specificData.transcriptomics:` Table with a column with Entrez ID’s and a column for the corresponding transcriptomics expression value in FPKM or raw counts (Default: empty).

| gene | expVal |
| --- | --- |
| entrezID1 | 220 |
| entrezID2 | 14916 |

**8**| (Optional) `XomicsToModel` can close the lower and upper bounds for ion exchange reactions, preventing net exchange of ions with the environment. To close the ions in the model, set `param.closeIons = true`. By doing so, it is possible to defer modelling of ion exchange (e.g., to represent action potentials in neurons), as they represent a dynamic trajectory of ion concentrations, while prioritising metabolic activities. Enabling ion exchange could be done where there is experimental evidence for net secretion or uptake of ions. The rate of uptake or secretion should then be specified for each ion in Step 14.

#### Usable Variable for Step 8

`param.closeIons:` Logical, true if ion exchange reactions are to be closed. (Default: false).

**▲ CRITICAL** If the parameter `param.closeUptakes` is set to `true`, ensure that uptakes are included; otherwise, it may lead to inconsistent models due to the absence of uptake reactions.

**9**| (Optional) To close the flux of exchange reactions, set `param.closeUptakes = true`. When this action is taken, the lower bound of the exchange reactions in the draft model are closed (set to 0) except those reactions subject to `specificData.rxns2constrain`. This will allow the draft model to only take up the metabolites specified by the bibliomics, growth media, or metabolomics data.

Set `param.closeUptakes = true` for biological systems derived from defined cell cultures.

Review the log; if `param.printLevel` is greater than zero, the new constraints will be printed along with statistics such as the size of the stoichiometric matrix, the number of exchange reactions closed, the number of exchange reactions in active reactions, and the number of exchange reactions in reactions to constrain based on manual literature curation.

#### Usable Variables for Step 9

`param.closeUptakes:` Logical, decide whether or not all of the uptakes in the draft model will be closed (Default: false).

`specificData.rxns2constrain:` Table containing the reaction identifier, the updated lower bound (lb), the updated upper bound (ub), a constraint description, and any notes such as references or special cases (Default: empty).

| rxns | lb | ub | constraintDescription | Notes |
| --- | --- | --- | --- | --- |
| rxn1 | 0 |  | Close uptake | PMID: **** |
| rxn2 | 0.0.956 | 1000 | Turnover constraints | PMID: **** |

### Close sink and demand reactions (optional)

**10**| (Optional) To set the demand and sink reactions to be closed, edit the parameter `param.nonCoreSinkDemands = true`. The demand and sink reactions specified in `specificData.activeReactions` will be exempt from closure. This distinction is vital, as closed reactions are not considered by the extraction algorithm for inclusion in the final model.

The solution of a feasible model must satisfy the constraints *Sv* = 0 and *l* ≤ *v* ≤ *u* and optionally coupling constraints *Cv* ≤ *d*. The newly added constraints may not all be consistent with a steady state flux at the same time, implying that the system of inequalities (and therefore the model) are infeasible, which could be caused by an incorrectly specified reaction bound. To perturb the constraints on an infeasible model to make it feasible, we use [relaxedFBA](https://opencobra.github.io/cobratoolbox/stable/modules/analysis/relaxedFBA/index.html?highlight=relaxedFBA#src.analysis.relaxedFBA.relaxedFBA), a constraint relaxation function that uses cardinality optimisation to find a minimal number of constraints to relax in order to make a model admit a steady state flux vector (i.e., a feasible Flux Balance Analysis problem). Cardinality optimisation is implemented by approximating it with an efficient continuous optimisation algorithm [11].

Set the relaxation options in `param.relaxOptions` (e.g., to only relax bound constraints or steady state constraints) for all or defined sets of reactions or metabolites.

`XomicsToModel` saves the old bounds from spent media metabolites in two new fields in the draft model model (`model.lbpreSinkDemandOff` and `model.ubpreSinkDemandOff`), along with fields identifying the reactions that were relaxed (see [relaxedFBA](https://opencobra.github.io/cobratoolbox/stable/modules/analysis/relaxedFBA/index.html?highlight=relaxedFBA#src.analysis.relaxedFBA.relaxedFBA)). `relaxedFBA` is a flexible function for relaxing constraints in `XomicsToModel.` The user may control it by setting the parameters in `param.relaxOptions` (e.g., to relax steady state, flux bounds or subsets and combinations thereof). Possible options are described in detail in Step 23 of COBRA Toolbox v3.0 protocol [38].

Review the log; if `param.printLevel` is greater than zero, the number and names of closed demand or sink reactions are printed.

#### Usable Variables for Step 10

`param.nonCoreSinkDemands:` The type of sink or demand reaction to close is indicated by a string (Possible options: `‘closeReversible’`, `‘closeForward’`, `‘closeAll’`, and `‘closeNone’`; Default: `‘closeNone’`).

`param.relaxOptions:` A structure array with the relaxation options; Default: `param.relaxOptions.steadyStateRelax = 0`). `specificData.activeReactions:` List of reactions known to be active based on bibliomic data (Default: empty).

### Set metabolic constraints (optional)

**CRITICAL** Extracellular metabolic constraints can be obtained from two different sources: fresh cell culture media metabolite concentrations (as specified by the manufacturer, or as specified by combining known quantities of metabolically specified media), or quantitative extracellular metabolomic measurements of fresh and spent media (assuming a metabolomic steady state), or both. The difference between spent and fresh media concentrations for extracellular metabolites is assumed to equal the rate of exchange of the metabolite between the cells and cell culture media (discussed further below). Based on your overall objective, constraints from fresh cell culture media metabolite concentrations or extracellular metabolomic measurements can either be integrated before or after the extraction algorithm step.

**11**| (Optional) To integrate constraints, set the `param.growthMediaBeforeModelExtraction` and `param.metabolomisBeforeModelExtraction` parameters respectively. Consider the following important factors when specifying the set of reactions to extract.

- The default is `param.growthMediaBeforeModelExtraction = true` and `param.metabolomisBeforeModelExtraction = true` as it helps to ensure that the model that enters (and leaves) the extraction step is consistent with metabolomic data.
- Setting `param.growthMediaBeforeModelExtraction = false` and `param.metabolomisBeforeModelExtraction = false` usually results in a less constrained model, which can be useful if one wishes to change the exometabolomic constraints (e.g., to model perturbations). However, care must be taken to ensure reactions that exchange metabolites corresponding to exometabolomic measurements are still present in the extracted model.

If the constraints are added before the extraction, the draft model solution space will be smaller. If it is done after the extraction, the context-specific model is extracted from a larger solution space with only the reaction directionality, steady-state and perhaps coupling constraints, given a particular network input into the model extraction step.

#### Usable Variables for Step 11

`param.growthMediaBeforeModelExtraction:` Logical, should the cell culture data be added before (true) or after (false) the model extraction (Default: true). `param.metabolomisBeforeModelExtraction:` Logical, should the metabolomic data be added before (true) or after (false) the model extraction (Default: true).

### Cell culture data (optional)

**CRITICAL** The uptake rates for exchange reactions corresponding to metabolites present in the media are adjusted by `XomicsToModel` based on their concentration by modifying the lower bounds on uptake rates, assuming the maximum uptake rate completely removes each metabolite from the medium between media replacements.

**12**| (Optional) Use the function supplementary function `preprocessingOmicsModel`, described in the Supplementary Information, to calculate the maximum medium uptake rates based on the fresh media metabolite concentrations. Set the field `mediumConcentrations` in `specificData.mediaData` to incorporate either the measured concentrations of metabolites in fresh media or those obtained from specifications of the media components. This will include the fresh media metabolite concentrations in subsequent calculations.

#### Usable Variable for Step 12

`specificData.`mediaData: Table containing the fresh media concentrations. Contains the reaction identifier, the maximum uptake (μ mol/gDW/h) assuming it is equal to the concentration of the metabolite in fresh medium divided by the length of time before medium exchange, and the medium concentration (μ mol/l; Default: empty).

| rxns | mediumMaxUptake | mediumConcentrations |
| --- | --- | --- |
| EX_met1 | -220 | 1345 |
| EX_met2 | -14916 | 191021 |

### Metabolomics (optional)

**▲ CRITICAL** Metabolomic exchange reaction constraints can be derived from quantitative metabolomic experiments. In the context of `XomicsToModel`, it is assumed that these measurements originate from the same biological sample at two different time-points Assess the quality of the data carefully before integrating it with the model by following the general community standards (measurement accuracy, precision, blank effect, linearity of the calibration line, relative errors of calibration line fit etc.) [54].

**▲ CRITICAL** The measured metabolites should be mapped onto the model namespace where metabolite identifiers and/or exchange reaction identifiers for each metabolite should be found (ideally by cross-mapping standard metabolite identifiers such as InChI, ChEBI, HMDB etc.) and included in the final metabolomics input data.

**13**| (Optional) Specify the rates of consumption or production based on exometabolomic data in `specificData.exoMet`. Outlier samples should be identified (for example, based on the median and quartiles of detected metabolite levels) and excluded. The metabolomics data is transformed into metabolic rates and normalised for the biomass content, e.g., per gram of dry weight (*gDW*) with:

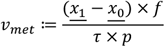

where *v* met is the rate of exchange of a given metabolite, *x*_0_ is the mean initial metabolite concentration (e.g., in *μmol*/*L*), *x*_1_ is the mean metabolite concentration at the second time point (*μmol*/*L*), *τ* is the time interval between the two measurements (in hours), *p* is the protein concentration in the sample (*g*/*L*), and *f* is the protein fraction of the total dry weight (g protein/g DW). It should be noted that the biomass content can also be estimated based on the cell count and an estimate of the dry weight mass of a cell or by direct measurements of dry weight mass (tissue). To propagate the error associated with exchange flux measurement, it is possible to propagate the standard deviation associated with each measured value using:

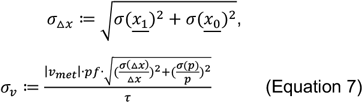

where *σ*(*Δx*) is the standard deviation in the estimation of the change in metabolite concentration between initial (*x*_0_) and final measurement (*x*_1_) given the measurement uncertainties, *σ*(*x*_1_) and *σ*(*x*_2_) respectively, and *σ*_*v*_ is the standard deviation in the exchange reaction rate, taking into account the standard deviation in metabolite concentration change estimation *σ*(*Δx*) and standard deviation in protein estimation *σ*(*p*). As the standard deviation in protein estimation is the same for each metabolite, when fitting a model that is linear in the rate of each reaction, one can consider this to be zero. The metabolomic constraints are set by fitting the bounds of the draft model to the exometabolomic reaction rates while allowing relaxation of net flux bounds based on the quadratic optimisation problem presented in previous work [9]:

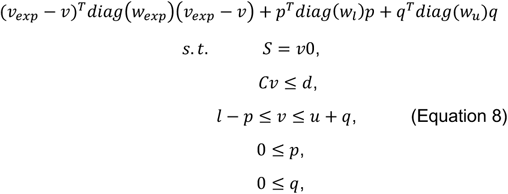

where 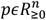and 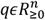are non-negative variables that allow the lower and upper bound constraints to be relaxed, *l* and *u*, respectively. The matrix-vector inequality *Cv* ≤ *d* represents coupling constraints, where the data *CϵR*^*s*×*n*^ and *dϵR*^*s*^ enforce *s* coupling constraints between net reaction fluxes (e.g., to represent a constraint on cumulative turnover of a metabolite by a set of degradation reactions). This problem always returns a steady-state flux *vϵR*^*n*^ and allows for different information to be input as parameters, including the penalisation of deviation from experimental fluxes 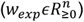, penalizing relaxation of lower bounds 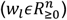, and penalising relaxation of upper bounds 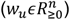.

Define the weights for penalizing the experimental fluxes in `param.metabolomicWeights`. By default, these weights are derived from the `‘mean’` of the experimental reaction flux. The weights could be derived from the standard deviation (*σv* in (2), ‘`SD`’) or relative standard deviation (‘`RSD`’) of the experimental reaction flux. For example, `param.metabolomicWeights =` ‘`SD`’ sets the penalty on deviation from experimental measurement to be the inverse of one plus the variance:

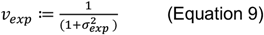

where 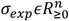is the standard deviation in measured fluxes. This approach increases the penalty on deviation from an experimentally measured mean flux where the variance is lower.

To avoid numerical errors in model analysis, a default flux value is used if the experimental data exceeds the specified boundary limits: the minimum reaction rate (`param.TolMinBoundary`), the maximum reaction rate (`param.TolMaxBoundary`) as well as the absolute minimum flux allowed, as defined in `param.boundPrecisionLimit`, i.e., if any |*l*_*j*_| or |*u*_*j*_| < `param.boundPrecisionLimit`, then it is considered as zero.

Review log, if `param.printLevel` is greater than zero, the new metabolomically derived bounds on reaction fluxes are printed.

Finally, the feasibility of the draft model is tested, and if it is not feasible, e.g., if there are inconsistencies or errors in the exometabolomic data, `XomicsToModel` will relax some reaction bounds using `relaxedFBA`.

#### Usable Variables for Step 13

`param.boundPrecisionLimit:` Precision of flux estimate, if the absolute value of the lower bound or the upper bound is lower than the *boundPrecisionLimit* but higher than 0 the value will be set to the *boundPrecisionLimit* (Default: primal LP feasibility tolerance).

`>> feasTol = getCobraSolverParams(‘LP’, ‘feasTol’);`

`>> param.boundPrecisionLimit = feasTol*10; param.metabolomicWeights:` String indicating the type of weights to be applied for metabolomics fitting (Possible options: `‘SD’`, `‘mean’` and ‘RSD’; Default: `‘mean’`). `param.TolMaxBoundary:` The reaction flux upper bound maximum value (Default: 1000).

`param.TolMinBoundary:` The reaction flux lower bound minimum value (Default: −1000). `specificData.essentialAA:` List exchange reactions of essential amino acids (Default: empty). Must never be secreted, even in a relaxed FBA model.

`specificData.exomet:` Table with measured exchange fluxes, e.g., obtained from an exometabolomic analysis of fresh and spent media. It includes the reaction identifier, the reaction name, the measured mean flux, standard deviation of the measured flux, the flux units, and the platform used to measure it (Default: empty).

| rxns | rxnNames | mean | SD | units | platform |
| --- | --- | --- | --- | --- | --- |
| EX_mMet1 | Exchange of mMet1 | 100 | 8x10<sup>-4</sup> | μ mol/gDW/h | LC-MS |
| EX_mMet2 | Exchange of mMet2 | 4x10<sup>-3</sup> | 3x10<sup>-4</sup> | μ mol/gDW/h | GC-MS |

### Add custom constraints (optional)

**CRITICAL** A custom constraint enables the specification of precise bounds on reactions, derived from bibliomic data, or the enforcement of essential reaction activity within the network.

**14**| (Optional) Specify bibliomic constraints in the struct array `specificData.rxns2constrain` described in Usable Variables for Step 14 to add both internal and external constraints on reactions. This information will be used to change the lower and upper bounds of the reactions in the draft model. Demand reactions will be ignored to ensure thermodynamic consistency if ‘`thermoKernel’` is chosen as the tissue-specific solver. This means that constraints on demand reactions are not considered, which helps maintain the thermodynamic consistency of the model generation process. No change is made if the bound is empty. If the bounds in the table exceed the established limit bounds, they will be adjusted to what the user or default data specifies. Finally, the feasibility of the draft model is tested and, if it is not feasible, `XomicsToModel` will relax some reaction bounds using [relaxedFBA](https://opencobra.github.io/cobratoolbox/stable/modules/analysis/relaxedFBA/index.html?highlight=relaxedFBA#src.analysis.relaxedFBA.relaxedFBA).

Review the log; if `param.printLevel` is greater than zero, the custom constraints with non-default bounds, will be printed. If any discrepancies or issues are identified, refer to the troubleshooting section to resolve common constraint errors (e.g., bounds exceeding limits).

#### ? TROUBLESHOOTING

##### Usable Variables for Step 14

`param.boundPrecisionLimit:` Precision of flux estimate, if the absolute value of the lower bound or the upper bound is lower than the *boundPrecisionLimit* but higher than 0 the value will be set to the *boundPrecisionLimit* (Default: primal LP feasibility tolerance).

`>> feasTol = getCobraSolverParams(‘LP’, ‘feasTol’);`

`>> param.boundPrecisionLimit = feasTol*10; param.tissueSpecificSolver:` The name of the tissue-specific solver to be used to extract the context-specific model (Possible options: `‘thermoKernel’` and `‘fastcore’`; Default: `‘thermoKernel’`).

`param.TolMaxBoundary:` The reaction flux upper bound maximum value (Default: 1000).

`param.TolMinBoundary:` The reaction flux lower bound minimum value (Default: −1000). `specificData.rxns2constrain:` Table containing the reaction identifier, the updated lower bound (lb), the updated upper bound (ub), a constraint description, and any notes such as references or special cases (Default: empty).

| rxns | lb | ub | constraintDescription | Notes |
| --- | --- | --- | --- | --- |
| rxn1 | 0 |  | Close uptake | PMID: **** |
| rxn2 | 0.0.956 | 1000 | Turnover constraints | PMID: **** |

### Set coupled reactions (optional)

**▴ CRITICAL** If `param.addCoupledRxns` and the table `specificData.coupledRxns` are empty, this step is not performed.

**▲ CRITICAL** Often, published biochemical data specify metabolites that are known to be produced or metabolised by the biological system. However, a model frequently includes multiple production or degradation pathways. To ensure that the net production or consumption of the metabolite is represented correctly in the model, the concept of coupled reactions can be used [35]. In non-growing non-dividing cells, such as neurons, these constraints can be used to replace the biomass reaction to set constraints that specify specific biochemical requirements for cell maintenance [9].

**15**| (Optional) Unless using default settings, set the coupled reactions on `specificData.coupledRxns`. Coupling constraints are denoted by the matrix inequality *Cv* ≤ *d*, where the matrix *C* ∈ {−1,1}^*k*×*n*^ specifies whether a metabolite is produced or consumed and *d* ∈ *R*^*k*^ specifies the net value of the production or consumption of that metabolite. That is, *C*_*ij*_ = −1 if metabolite *i* is consumed by reaction *j* and *C*_*ij*_ = 1 if metabolite *i* is produced by reaction *j*. This constraint ensures that some of the coupled reactions are active in the model and the total production or consumption is at least the amount given by *d*_*i*_. However, the exact flux partition of total flux within the set of reactions is not specified. The coupling constraints are added to the draft model by including fields such as the constraint matrix containing reaction coefficients `model.c`, the constraint identifiers `model.ctrs`, the constraint senses `model.dsense`, and the constraint right-hand side values `model.d`.

Review the log; if `param.printLevel` is greater than zero, the information about the added coupled reactions will be printed. For the *i*th coupling constraint, when `model.dsense(i) = ‘G’` this implies:

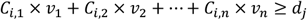

and when `model.dsense(i) = ‘L’` it implies:

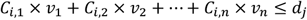

Within the pipeline, the function `addCOBRAConstraints` adds the specified set of coupling constraints to the model.

#### ? TROUBLESHOOTING

##### Usable Variables for Step 15

`param.addCoupledRxns:` Logical, should the coupled constraints be added (Default: false).

`specificData.coupledRxns:` Table containing information about the coupled reactions. This includes the identifier for the coupling constraint (couplingConstraintID), the list of coupled reactions (coupledRxnsList), the coefficients of those reactions (c, given in the same order), the right-hand side of the constraint (d), the constraint sense or the directionality of the constraint (dsence), and the reference (Default: empty).

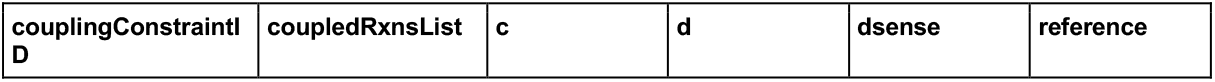
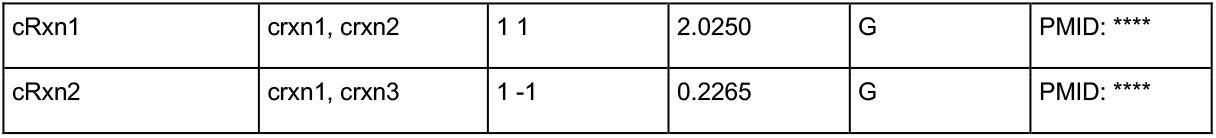

| couplingConstraintID | coupledRxnsList | c | d | dsense | reference |
| --- | --- | --- | --- | --- | --- |
| cRxn1 | crxn1, crxn2 | 1 1 | 2.0250 | G | PMID: **** |
| cRxn2 | crxn1, crxn3 | 1 -1 | 0.2265 | G | PMID: **** |

### Remove inactive reactions

**▴ CRITICAL** Prior to the extraction process, it is crucial to selectively exclude certain reactions from the draft model emphasizing the importance of refining the model’s accuracy and relevance. This process ensures that subsequent computational analyses are conducted on a streamlined and biologically meaningful metabolic network.

**16**| (Optional) Identify and list metabolic reactions that should be inactive by manual curation of the literature (`specificData.inactiveReactions`). Designate reactions for removal from the draft model, considering them non-essential (`specificData.rxns2remove.rxns`). This encompasses reactions such as sink or demand reactions that might not be deemed vital for the model. In the event of a discrepancy between input data (e.g. a reaction assigned to be inactive based on manual literature curation but active according to Step 5), specify the preference between curated data or omics data from experiments using the parameter `param.curationOverOmics`.

- Set `param.curationOverOmics = true` to prioritise manual literature curation
- Set `param.curationOverOmics = false` to prioritise experimental data.

After reaction removal, the feasibility of the model is tested and, if it fails, some reaction bounds will be relaxed via [relaxedFBA](https://opencobra.github.io/cobratoolbox/stable/modules/analysis/relaxedFBA/index.html?highlight=relaxedFBA#src.analysis.relaxedFBA.relaxedFBA).

Review the log; if `param.printLevel` is greater than zero, the number of reactions removed and the reactions that were assigned as inactive and were removed will be printed.

#### Usable Variables for Step 16

`param.curationOverOmics:` Logical, indicates whether curated data should take priority over omics data (Default: false).

`specificData.inactiveReactions:` List of reactions known to be inactive based on manual literature curation (Default: empty).

`specificData.rxns2remove.rxns:` List of reactions to remove from the generic model (Default: empty).

### Remove inactive genes

**▴ CRITICAL** Building on the previous step (Step 16), selectively removing certain genes from the draft model enhances both the quality and specificity of the metabolic network.

**17**| Identify and list metabolic genes known to be inactive based on the bibliomics data (`specificData.inactiveGenes`). Genes assigned as inactive in bibliomic or transcriptome data are removed from the draft model by `XomicsToModel`, together with the reactions affected by their deletion (identified using the function [allRxnPerActiveGene](https://opencobra.github.io/cobratoolbox/stable/modules/analysis/deletion/index.html?highlight=deleteModelGenes#src.analysis.deletion.deleteModelGenes)). However, any reaction assigned to be active in Step 5 will not be deleted at this step. Furthermore, if removing reactions that correspond to inactive genes compromises the feasibility of a draft model, the bounds of the fewest number of those reactions are relaxed via [relaxedFBA](https://opencobra.github.io/cobratoolbox/stable/modules/analysis/relaxedFBA/index.html?highlight=relaxedFBA#src.analysis.relaxedFBA.relaxedFBA), and the rest are removed. To ensure that the model is feasible, the relaxed reactions are added to the list of active reactions. In the event of a discrepancy between the datasets, for example, a gene assigned to be inactive based on the manual literature curation but active based on the proteomics data or transcriptiomics data, or both (Step 7), preference will be given based on the value of `param.curationOverOmics` variable.

Review the log; if `param.printLevel` is greater than zero, the deleted reactions and genes will be printed, as well as those that were not deleted.

#### Usable Variables for Step 17

`param.curationOverOmics:` Logical, indicates whether curated data should take priority over omics data (Default: false).

`specificData.inactiveGenes:` List of Entrez identifiers of genes known to be inactive based on the bibliomics data (Default: empty).

### Find flux consistent subset

**▴ CRITICAL** `XomicsToModel` finds and extracts the largest subset of the draft model that is flux consistent, i.e. each reaction that can carry a non-zero steady-state flux.

**18**| Unless using default settings, specify the parameters `param.fluxCCmethod` and `param.fluxEpsilon` to indicate the algorithm to be used to identify the flux consistent subset of the draft model and to determine the minimum flux magnitude considered non-zero. All reactions that are flux inconsistent and all metabolites that exclusively participate in flux inconsistent reactions are removed by `XomicsToModel`. If a flux inconsistent metabolite or reaction is identified as active in Step 5, `XomicsToModel` removes it from the list.

Review the log; if `param.printLevel` is greater than zero, `XomicsToModel` prints the dimensions of the stoichiometric matrix before and after checking flux consistency, as well as the constraints for flux consistent reactions and metabolites.

In addition, different boolean vectors indicating the flux consistent and inconsistent metabolites and reactions are added to the draft model by `XomicsToModel`.

#### Usable Variables for Step 18

`param.fluxCCmethod:` String with the name of the algorithm to be used for the flux consistency check (Possible options: `‘swiftcc’`, `‘fastcc’` or ‘dc’, Default: `‘fastcc’`). `param.fluxEpsilon:` Minimum non-zero flux value accepted for tolerance (Default: Primal feasibility tolerance).

`>> feasTol = getCobraSolverParams(‘LP’, ‘feasTol’);`

>> param.fluxEpsilon = feasTol * 10;

### Find thermodynamically consistent subset (optional)

**CRITICAL** In this step, `XomicsToModel` finds the largest thermodynamically consistent subset. The performance of `thermoKernel` is accelerated if provided with a thermodynamically consistent model prior to extracting a subset. `XomicsToModel` finds the largest thermodynamically consistent subset using `findThermoConsistentFluxSubset`, which requires `model.S`, `model.lb` and `model.ub` as essential inputs.

**19**| (Optional) Unless using default settings, specify `param.tissueSpecificSolver = ‘thermoKernel’` as the extraction algorithm. Specify the metabolites and reactions that are flux consistent using `model.fluxConsistentMetBool` and `model.fluxConsistentRxnBool` to accelerate this step.

All reactions that are not thermodynamically consistent are removed from the draft model by `XomicsToModel`. All metabolites that are exclusively involved in thermodynamically inconsistent reactions are also removed from the draft model by `XomicsToModel`. If a thermodynamically inconsistent metabolite or reaction is identified as active in `specificData.presentMetabolites` or in the list of active reactions from Step 5, it is removed from the list by `XomicsToModel`. Additionally, all orphan genes are removed from the model and the reaction-gene-matrix is regenerated. Boolean vectors indicating thermodynamically consistent metabolites and reactions are added to the draft model, `model.thermoFluxConsistentMetBool` and `model.thermoFluxConsistentRxnBool,` respectively.

Review the log; if `param.printLevel` is greater than zero, `XomicsToModel` prints the dimensions of the stoichiometric matrix before and after checking thermodynamic consistency, as well as the constraints for thermodynamic consistent reactions and metabolites.

To identify a thermodynamically feasible flux, examine the net flux vector where each non-zero entry indicates a sign opposite the change in chemical potential for the corresponding reaction. A model is deemed thermodynamically consistent if each of its reactions allows for a non-zero thermodynamically feasible flux, thereby meeting the criteria for consistency. This is outlined by the following mathematical definition:

#### Usable Variables for Step 19

`param.tissueSpecificSolver:` The name of the tissue-specific solver to be used to extract the context-specific model (Possible options: `‘thermoKernel’` and `‘fastcore’`; Default: `‘thermoKernel’`).

`param.thermoFluxEpislon:` Flux epsilon used in `thermoKernel` (Default: solver primal feasibility tolerance × 10).

`param.iterationMethod:` Enables different iteration methods to be employed when exploring the thermodynamically consistent set.

### Identify active reactions from genes

**▲ CRITICAL** The list of active reactions up to this point has been based on the subset of user-specified context-specific reactions that are determined to be flux consistent by `XomicsToModel`. The relationship between context-specific genes and metabolic reactions is established by `XomicsToModel` in this step. Each identified reaction is added by `XomicsToModel` to the final list of active reactions used to extract the context-specific model.

**20**| Decide how `XomicsToModel` determines the gene-reaction relationship and specify your preference using `param.activeGenesApproach`:

- `‘oneRxnPerActiveGene’`: At least one reaction per active gene is included (default).
- `‘allRxnPerActiveGene’`: Find a list of reactions, whose rates are affected by the deletion of active genes and include them all as active reactions.

Changing this parameter has a large effect on the size of the extracted models. Per active gene, `oneRxnPerActiveGene` adds at least one corresponding reaction to the extracted model, whereas `allRxnPerActiveGene` adds all corresponding reactions, where the presence of the gene product is essential for nonzero flux through each of the reactions corresponding. Experience with generation of whole-body metabolic models [39] and extraction of a neuronal model [9] from Recon3D supports the use of `oneRxnPerActiveGene` as the default.

#### Usable Variable for Step 20

`param.activeGenesApproach:` String with the name of the active genes approach will be used (Possible options; `‘oneRxnPerActiveGene’`, `‘allRxnPerActiveGene’`; Default: `‘oneRxnPerActiveGene’`).

### Model extraction

**▴ CRITICAL** In this step, given a draft model, a specific model is extracted using the function `createTissueSpecificModel`. The `createTissueSpecificModel` function is already enabled to extract context-specific models using a suite of model extraction algorithms (GIMMME [22], iMAT [23], MBA [7], INIT [55], mCADRE [24] and FASTCORE

[12]) and has been described previously (Step 40 of a previous protocol [38]). Each of these algorithms extracts a flux-consistent context-specific model. In `XomicsToModel`, the default model extraction algorithm is `thermoKernel`, because it enables extraction of a context-specific model that is stoichiometrically consistent, flux-consistent and thermodynamically consistent, now accessible via the `createTissueSpecificModel`. Implementation of thermodynamic constraints during the model extraction process enables integration of context-specific data on each metabolite, which is not directly integrable with the other aforementioned model extraction algorithms.

**21**| Set the `param.tissueSpecificSolver` parameter to guide on the choice of solver: The default option `‘thermoKernel’` extracts a thermodynamically consistent context-specific model, ensuring a robust foundation for further analyses. Alternatively, `‘fastCore’` for extracting a minimal set of reactions supporting optimal flux solutions, emphasizing the preservation of essential functionalities.

#### Usable Variable for Step 21

`param.`tissueSpecificSolver: The name of the tissue-specific solver to be used to extract the context-specific model (Possible options: ‘thermoKernel’ and ‘fastCore’; Default: ‘thermoKernel’).

### thermoKernel (optional)

**▴ CRITICAL** `thermoKernel` is an algorithm designed to extract a thermodynamically consistent context-specific model. It is important that the weightings on metabolites and weightings on reactions (derived from experimental data) are appropriately adjusted by the user to ensure accurate and context-specific model outcomes.

**22**| **(Optional)** To assign weights to metabolites, based on quantitative data or through manual curation, set the weights of the metabolites in `specificData.presentMetabolites.weights` accordingly (see Experimental Design). Likewise, for assigning weights to absent metabolites, set `specificData.absentMetabolites.weights` to values < 0. Negative weights promote inclusion of metabolites and reactions in a specific model, positive weights do the opposite, and a zero weight does not penalise for exclusion or inclusion.

By default, `XomicsToModel` sets a fixed penalty for inclusion of any metabolite or reaction not in a core set. An exception is that `XomicsToModel` sets a weight of zero for any metabolite ranked in the top 100 for metabolite connectivity, mostly non-specific cofactors. This is to avoid exclusion or inclusion metabolites, such as ATP, that participate in any context-specific network. Therefore, the objective of integrating metabolite weights is to promote exclusion or inclusion of metabolites that participate in a small number of relatively pathway-specific reactions.

#### Add a penalty for reactions

Provide gene expression data (Step 7) to set the fixed penalty equal to the median of the log-transformed reaction expression values in `model.expressionRxns`. In the absence of expression data, the fixed penalty is set to one. Use `param.activeGenesApproach = ‘oneRxnPerActiveGene’` (see Step 20), to specify that all metabolites and reactions in the core sets are assigned a weight equal to the *negative* of the median of the log of the reaction expression value `model.expressionRxns`, or −1 if `model.expressionRxns` is not provided.

If `model.expressionRxns` is provided by the user but a reaction is missing gene expression data, `XomicsToModel` assumes that the corresponding reaction has a weight of equal to the *negative* of the median of the log of the reaction expression values `model.expressionRxns`.

#### Context-specificity and thermodynamic consistency

A model is context-specific and thermodynamically flux consistent if it meets two criteria:

- Thermodynamic flux consistency: Each reaction admits a non-zero thermodynamically feasible flux.
- Context-specific enrichment: The model includes metabolites and reactions relevant to a particular context while excluding those that are not relevant to that context.

#### Evaluate the extracted model

An extracted model should be considered a draft model, until it has been compared with the expected results of model extraction. For example, if `thermoKernel` was selected as the model extraction algorithm, then all reactions should admit a thermodynamically feasible flux. Furthermore, the draft model should be checked to see if it contains the anticipated metabolites and reactions. If not, then adjustment of parameters or input data may be warranted.

Review the log; if `param.printLevel` is greater than zero, the size of the stoichiometric matrix, the metabolites and reactions removed by the extraction algorithm, and the core metabolites and reactions removed by the extraction solver are all printed.

- If greater than 1, the progress of each outer iterations (Equation 6) of the `thermoKernel` algorithm is printed, including the fraction of thermodynamically consistent reactions and metabolites that have been added to the extracted model at each iteration and their overlap with the incentivised and penalised reactions and metabolites.
- If greater than 2, the progress of each inner iteration (Equation 5) of the `thermoKernel` algorithm is printed, including the number of non-zero thermodynamically feasible fluxes identified in each iteration, before and after checking for thermodynamic feasibility, thus enabling the user to monitor the progress of the model extraction.

#### ? TROUBLESHOOTING

##### Usable Variables for Step 22

`param.activeGenesApproach:` String with the name of the active genes approach will be used (Possible options; `‘oneRxnPerActiveGene’`, `‘allRxnPerActiveGene’`; Default: `‘oneRxnPerActiveGene’`).

`param.fluxEpsilon:` Minimum non-zero flux value accepted for tolerance (Default: Primal feasibility tolerance).

`>> feasTol = getCobraSolverParams(‘LP’, ‘feasTol’);`

>> param.fluxEpsilon = feasTol * 10;

`param.thermoFluxEpsilon:` Minimal flux considered thermodynamically consistent by `thermoKernel` (Default: primal feasibility tolerance × 10). `param.tissueSpecificSolver:` The name of the tissue-specific solver to be used to extract the context-specific model (Possible options: `‘thermoKernel’` and `‘fastCore’`; Default: `‘thermoKernel’`).

`param.weightsFromOmics:` True for gene weights to be assigned based on the omics data (Default: true).

`specificData.absentMetabolites:` List of metabolites confirmed to be absent based on bibliomics data and their corresponding weights. `specificData.presentMetabolites:` List of metabolites confirmed to be present based on bibliomics data and their corresponding weights.

### Final adjustments

**CRITICAL** In the last step, `XomicsToModel` checks the flux and thermodynamic consistency of the model and adds any specific data and the parameters used in the extraction of the specific model to the model structure, that is `model.XomicsToModelSpecificData` and `model.XomicsToModelParam`, respectively. If a field with metabolic reaction formulas did not already exist in the draft model, `XomicsToModel` adds it using the `printRxnFormula` function [58]. Also, `XomicsToModel` adds two vectors that indicate which reactions were specified to be active or inactive (`model.activeInactiveRxn`), as well as which metabolites were specified to be present or absent (`model.presentAbsentMet`). Finally, `XomicsToModel` reorders the fields of the context-specific model into a standard format.

**23**| Review the log entries to confirm the model’s dimensions at various steps in the `XomicsToModel` process. For example, the effect of major steps in the pipeline are evident by the relative increase or decrease in size of the draft model, so unexpected changes in model size should be investigated in the corresponding step to ensure that the input data is correctly specified and the technical parameters are appropriate to the context.

If `param.printLevel` was set to > 0, the algorithm will display all outputs from the `XomicsToModel` function. This detailed output can be useful, especially in identifying steps where reactions were removed due to lack of flux consistency, stoichiometric consistency, or thermodynamic consistency. Such insights serve as cues for potential manual curation, allowing users to address specific reactions that might have been excluded but could be of interest. Additionally, having a detailed log facilitates traceability, enabling users to understand the evolution of the model throughout the process. It serves as a documentation tool, aiding in debugging, and provides a comprehensive record of the decisions made during model construction, offering transparency and reproducibility in the analysis.

#### TIMING

Step 1, Data Preparation: *∼* 5s

Step 2, Generic model check: *∼* 1s

Step 3, Set ob*j*ective function: 1s

Step 4, Add missing reactions: *∼* 30s

Step 5, Identify core metabolites and reactions: *∼* 10s

Step 6, Set limit bounds: *∼* 1s

Step 7, Identify active genes: *∼* 10^2^s

Step 8, Close ions: *∼* 1s

Step 9, Close exchange reactions: *∼* 1s

Step 10, Close sink and demand reactions: *∼* 10s

Step 11, Set metabolic constraints: *∼* 10s

Step 12, Cell culture data: *∼* 10s

Step 13, Metabolomics: *∼* 10s

Step 14, Add custom constraints: *∼* 10s

Step 15, Set coupled reactions: 10s

Step 16, Remove inactive reactions: 10s

Step 17, Remove inactive genes: *∼* 10^2^s

Step 18, Find flux consistent subset: *∼* 10^2^s

Step 19, Find thermodynamically consistent subset: *∼* 10s

Step 20, Identify active reactions from genes: *∼* 10s

Step 21, Model extraction: *∼* 10^2^s

Step 22, thermoKernel: *∼* 10^2^s

Step 23, Final adjustments: *∼* 1s

#### ? TROUBLESHOOTING

Troubleshooting advice can be found in Table 2.

**Table 2:**
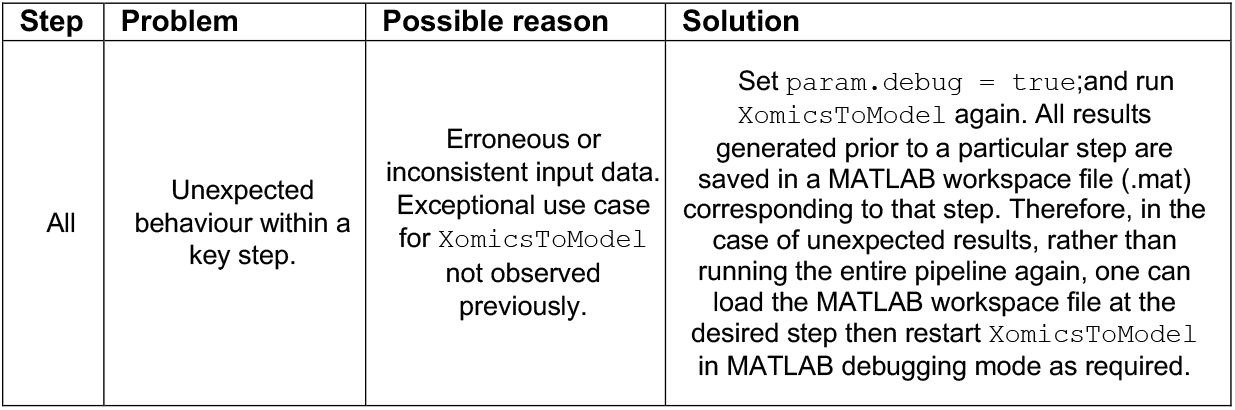
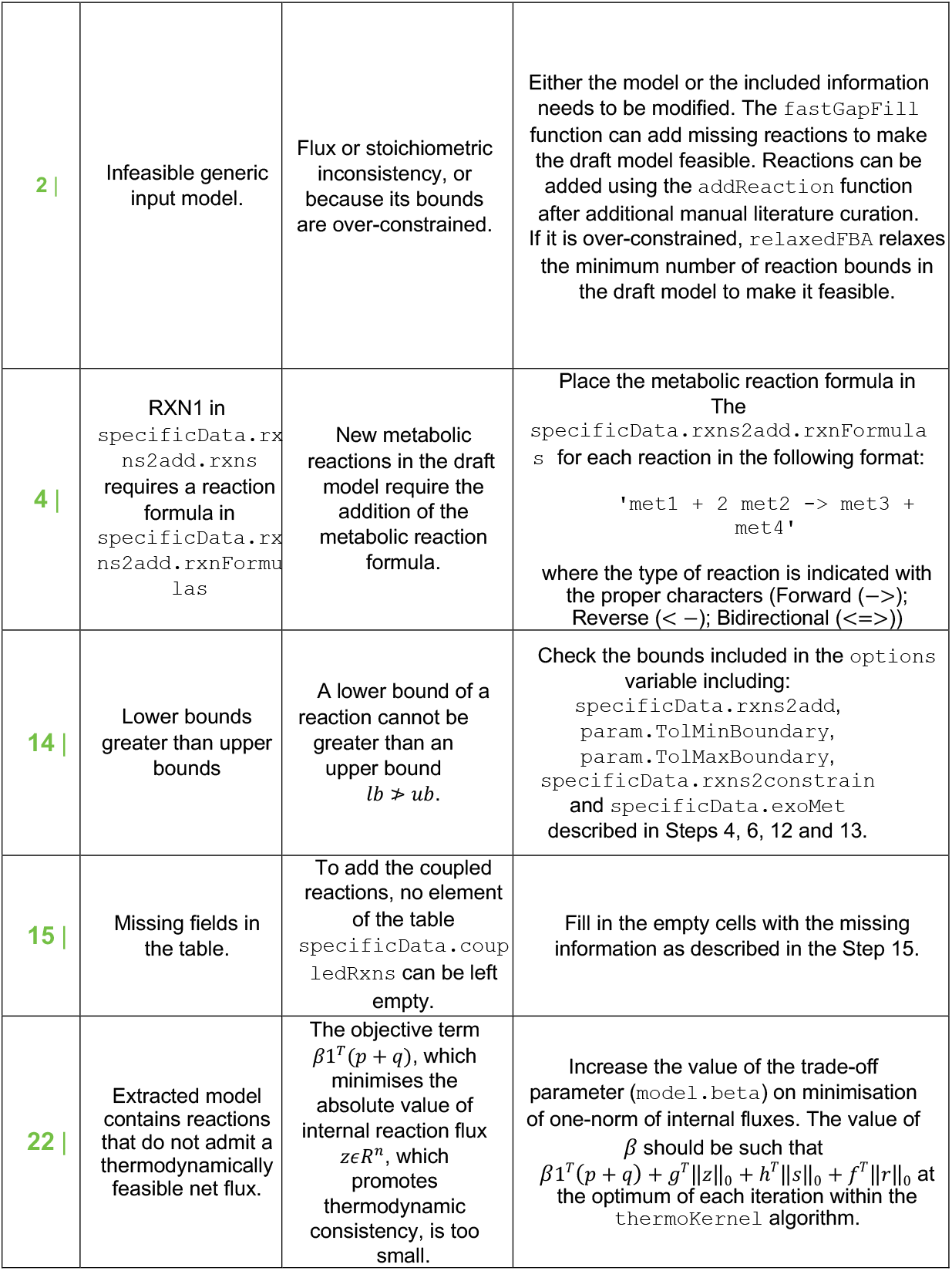
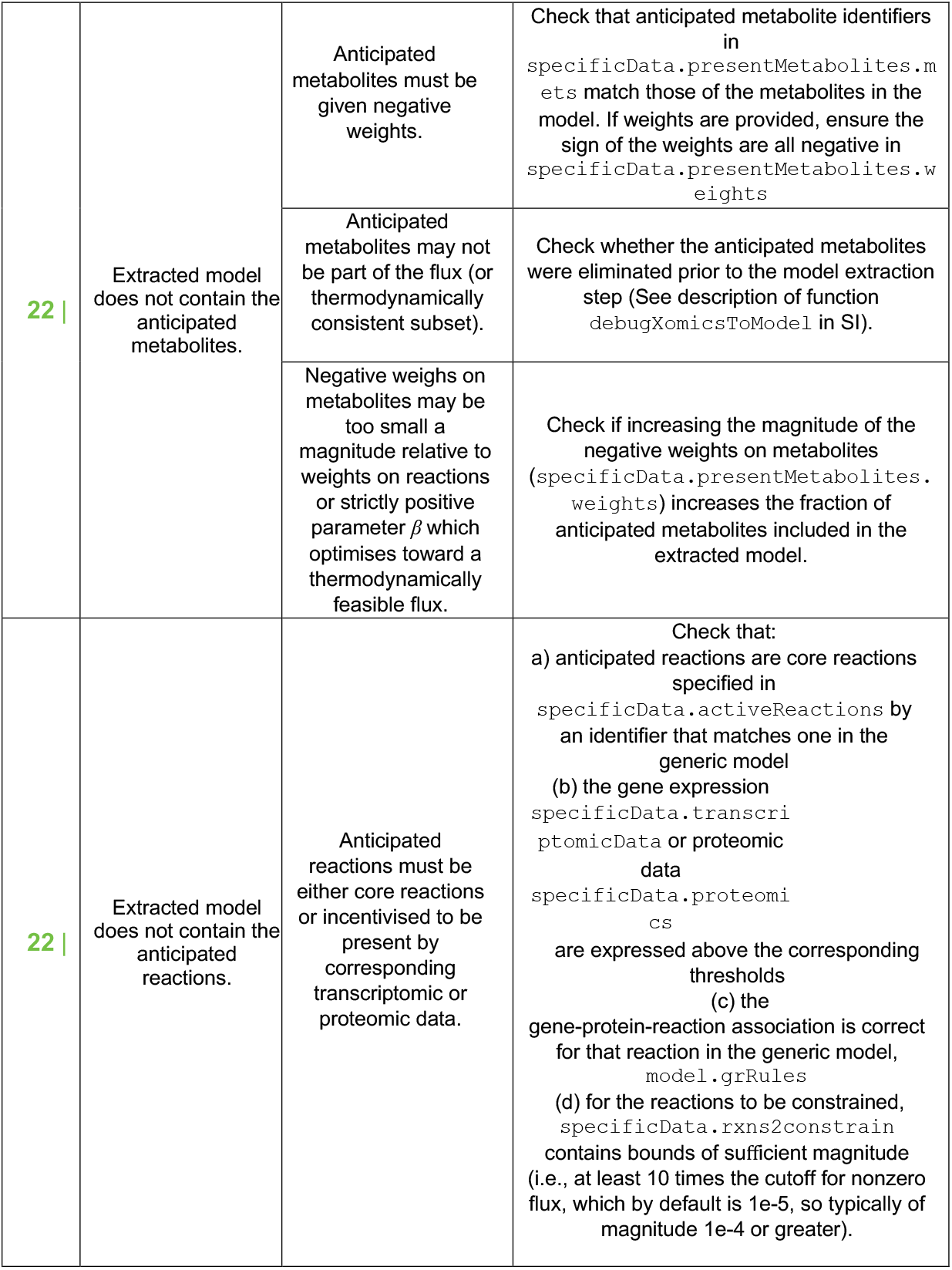
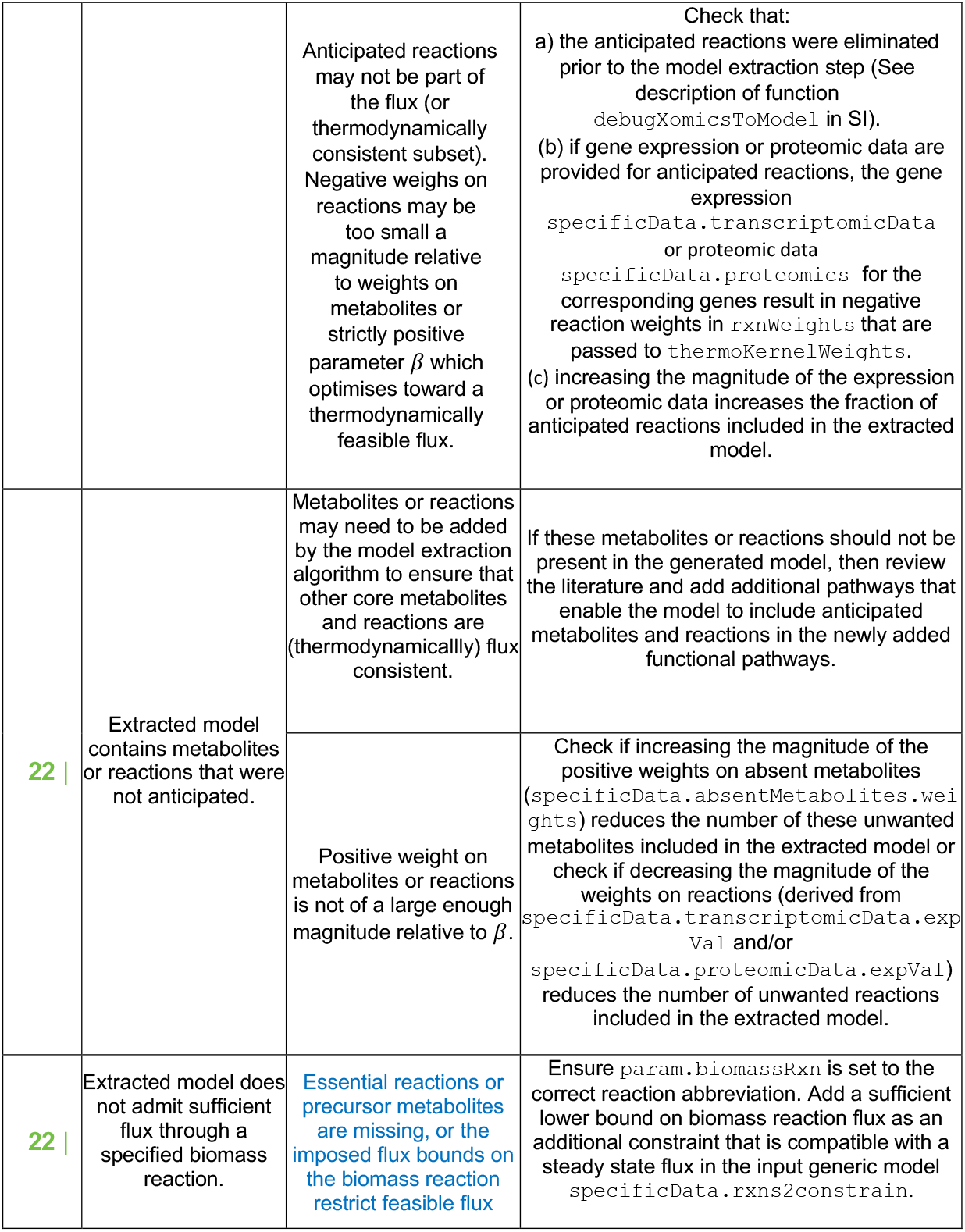
Troubleshooting.

**Table 3:** Recon3D [25] stoichiometric consistency summary.

| Mets | Rxns | Analysis |
| --- | --- | --- |
| 5835 | 10600 | totals |
| 0 | 1809 | heuristically external |
| 5835 | 8791 | heuristically internal |
| 5835 | 8791 | ... of which are stoichiometrically consistent |
| 0 | 0 | ... of which are stoichiometrically inconsistent. |
| 0 | 0 | ... of which are of unknown consistency. |
| 0 | 0 | heuristically internal and stoichiometrically inconsistent or unknown consistency. |
| 0 | 0 | ... of which are elementally imbalanced (inclusively involved metabolite). |
| 0 | 0 | ... of which are elementally imbalanced (exclusively involved metabolite). |
| 5835 | 8791 | Confirmed stoichiometrically consistent by leak/siphon testing. |

## ANTICIPATED RESULTS

This section follows the steps of the Procedure. Parameters used and modified are printed in the diary described in Step 2. Any results or anomalies are also printed in this diary.

**1**| The parameters that will be used by the `XomicsToModel` pipeline are printed, including the default values for options that have not been declared.

**2**| Any modifications made at this step are indicated; for example, if the generic model contains DM_atp_c_ reaction it is re-named to ATPM as it represents ATP **m**aintenance and to ensure that it is not closed together with standard demand reactions for the model extraction with `thermoKernel`. Furthermore, if the compartment representing mitochondrial intramembrane space ([i]) is removed (if `thermoKernel` is chosen) the modified reactions are also printed out.

**3**| The output of this set is a message including the specified linear objective function, or a lack of thereof.

**4**| The reactions and formulas added in this step are printed, together with the results of the stoichiometric consistency test. An example of the stoichiometric consistency test for Recon3D [25] can be seen in Table 3.

**5**| The output of this set is a set of present metabolites and reactions identified based on bibliomic, metabolomic, and cell culture data.

**6**| The output of this set is a draft model with the updated maximum upper bounds and minimum lower bounds.

**7**| The output of this set is a set of active genes identified using bibliomic, transcriptomic, and proteomic data. In addition, a message is displayed indicating the number of genes that have no expression information due to a lack of transcriptomic data.

**8**| The output of this set is a draft model in which the upper and lower bounds of ion exchange reactions such as calcium, potassium, or sodium, among others, are set to zero.

**9**| The output of this set is a draft model where all the lower bounds of exchange reactions are set to zero, except for those exchange reactions corresponding to metabolites specified to be present in the environment, e.g.,fresh media metabolites, which are assumed to be possible to be taken up from the media. Exchange reaction upper bounds are unchanged. A message is shown reporting model statistics such as the stoichiometric matrix dimensions, the number of closed reactions, the number of all exchange reactions, and the number of exchange reactions in the active reactions.

**10**| The sink and demand reactions of the draft model are set to zero and a message is shown reporting the number of closed non-core sink/demand reactions and the method used in param.nonCoreSinksDemands.

**11, 12 and 13**| The output of this set is a draft model with added metabolomic constraints from the environment, e.g., cell culture exometabolomic constraints, or both. Compared to the draft model from the previous step which usually derives generic upper and lower bounds from the generic input model, the addition of metabolomic constraints from the environment results in a subset of exchange reactions with quantitatively specified upper and lower bounds. In addition, a message is displayed reporting the number of exometabolomic and media uptake constraints. It includes exometabolomic data statistics, the fitting procedure for each exometabolomic reaction, the relaxed reactions and the summary of the fitting.

**14**| The output of this set is a draft model with custom constraints based on bibliomic data. If thermoKernel is being used, demand reactions included in specificData.rxns2constrain, i.e. with prefix DM_, will be ignored and the message indicating this is being printed. Furthermore, a message is given showing a list of reaction identifiers that cannot be constrained because they are not present in the model, the number of constrained reactions, and whether or not the model was still feasible after the addition of custom constraints.

**15**| The output of this set is a draft model that includes new fields that correspond to the coupled reactions. A message is shown including the details of the coupled constraints imposed on the draft model.

**16**| The output of this set is a draft model, with inactive reactions removed (based on bibliomic data). A message including the number of reactions that were removed from the model, a list of reactions that were not present in the draft model prior to the reaction removal, as well as the results of the model feasibility check.

**17**| The output of this set is a draft model with inactive genes removed and a detailed description of the number of genes that were removed or missing in draft model prior to this step. Based on the value of param.curationOverOmics a message is given about the genes that were kept or removed from the manually curated set of inactive genes due to the conflict with the provided omics data (measured as active in transcriptomics and/or proteomics). Furthermore, information is given about the number of genes that were not removed since their removal would lead to an infeasible model. Last, model feasibility check results are given, including any reaction that was relaxed if the model was not feasible.

**18**| The output of this set is a draft model with flux consistent reactions and metabolites is printed, along with a report containing model statistics such as stoichiometric matrix dimensions, flux consistent and inconsistent reactions and metabolites.

**19**| The output of this set is a thermodynamically consistent draft model. Additionally, a message is given with the parameters used to identify the thermodynamically consistent subset. It is followed by a detailed report from the cardinality optimisation. Last, the stoichiometric matrix dimensions, thermodynamically consistent and inconsistent reactions, and metabolites are printed out.

**20**| The output of this set is a set of reactions that must be active based on the active genes.

**21 and 22**| The output of this set is an extracted genome-scale metabolic model.

- If `fastCore` is selected as the extraction algorithm, a message is printed out with the extraction summary and the size of the new model.
- If `thermoKernel` is selected as the extraction algorithm, a message is printed out with the parameters used in the `thermoKernel` function, as well as `optCardThermo` and `optimizeCardinality` functions called by the `thermoKernel` function, an extraction summary, and statistics for the new model.

Two confusion matrices are generated that compare specified and actual metabolites and reactions in an extracted model (Figure 4). This comparison validates the accuracy and completeness of the extraction process, ensuring that the model faithfully represents the biological system. Accurate validation is crucial for reliable analyses and simulations based on the model. A figure with comparison of weights on active and inactive reactions and metabolites is also generated (Figure 5). This is useful for assessing if the relative weights on present and absent metabolites, as well as active and inactive reactions, passed as parameters to `thermoKernel`, are sufficient to incentivise and disincentivise the presence or absence of metabolites and reactions in the generated model. Since the generated model is a compromise between the weights on metabolites and reactions one may find that, some metabolites and reactions may be omitted at the expense of inclusion of other reactions and metabolites, respectively, depending on the input weights to `thermoKernel`. The user can modify the weights in `param.weightsFromOmics`, `presentMetabolites.weights` or `absentMetabolites.weights` to control the inclusion or exclusion of specific metabolites and reactions.

**Figure 4:**
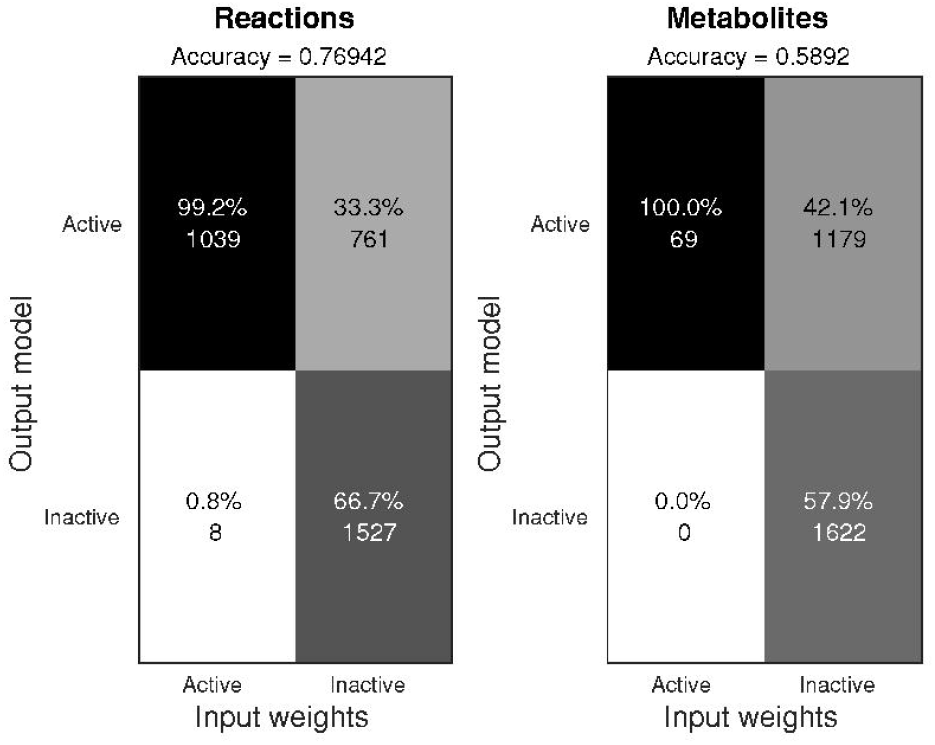
Comparison of specified and actual metabolites and reactions in an extracted model. Confusion matrices between the reactions and metabolites specified to be active and inactive versus output model reactions and metabolites that are actually active and inactive, obtained while generating a context-specific model of a dopaminergic neuronal metabolism [9]. The end result may be a trade-off between incentives to be active (negative weights), incentives to be inactive (positive weights) and ambivalence (zero weights). When a reaction is incentivised to be present but the corresponding metabolites are incentivised to be inactive, the optimal solution at each iteration of `thermoKernel` will depend on the relative weights on the reactions versus metabolites. This plot is automatically generated by `XomicsToModel` if `param.plotThermoKernelStats = 1;` when `param.tissueSpecificSolver = ‘thermoKernel’`.

**Figure 5:**
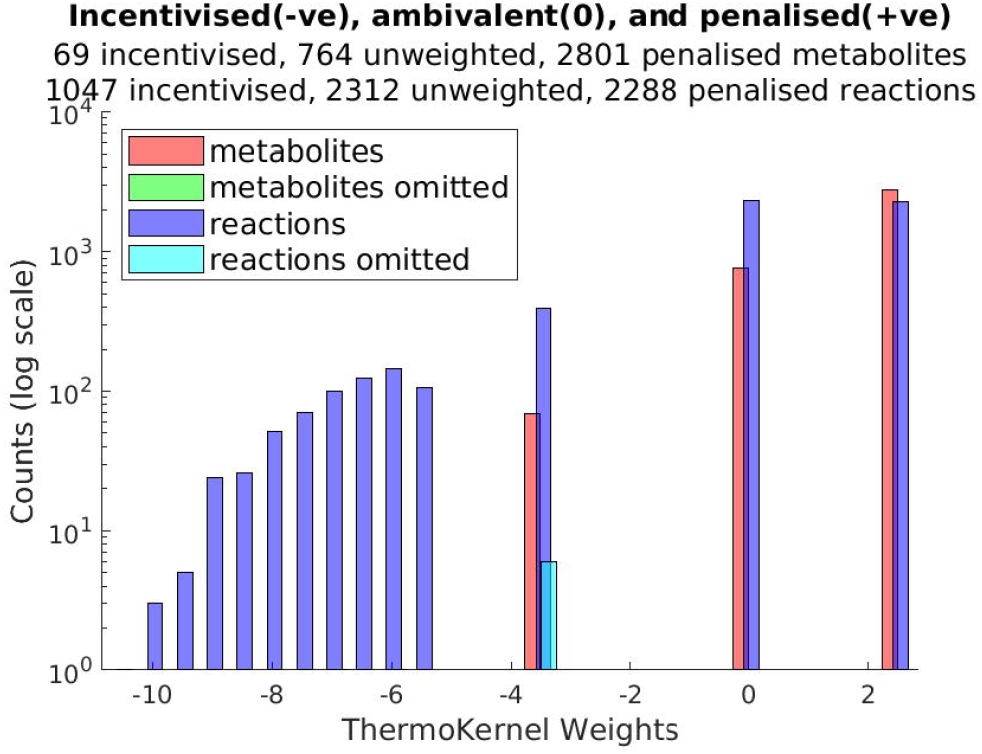
Comparison of weights on active and inactive reactions and metabolites. Almost all reactions incentivised to be active (negative weights) are active, with a small proportion inactive that should be active (reactions omitted, light blue), obtained while generating a context-specific model of a dopaminergic neuronal metabolism [9]. All metabolites incentivised to be active (negative weights) are active, with a none inactive that should be active (metabolites omitted, green in legend, no bar in plot). This plot is automatically generated by `XomicsToModel` if `param.plotThermoKernelStats = 1`; when `param.tissueSpecificSolver = ‘thermoKernel’`.

**23**| **Final adjustments** The ouput of this step is the final context-specific flux consistent genome-scale metabolic model. If `‘thermoKernel’` was chosen, then the model is also thermodynamically flux consistent. With all input data fully prepared, algorithmic completion of the pipeline takes ∼10 min, however manual review of intermediate results may also be required, e.g., when inconsistent input data lead to an infeasible model.

Previous applications of the `XomicsToModel` pipeline illustrate the type of results that can be anticipated. For instance, dopaminergic neuronal models generated with the protocol [9, 26] reproduced experimentally observed metabolite exchange rates and highlighted bioenergetic vulnerabilities relevant to Parkinson’s disease. Multi-omics integration studies [27, 28] demonstrated that `XomicsToModel` yields reproducible, condition-specific models that capture context-dependent metabolic rewiring. More recently, `XomicsToModel` was used to generate macrophage-specific models in Gaucher disease, a lysosomal storage disorder caused by *GBA1* mutations. These models integrated transcriptomic, exometabolomic, and bibliomic data to capture disease-specific perturbations, revealing mitochondrial dysfunction, a metabolic shift toward glycolysis, and profound alterations in lipid and cholesterol metabolism [29]. Furthermore, its use in isotope-labelled flux studies [30, 31] shows the potential of the protocol to provide mechanistic insight into immune metabolism and quantitative flux predictions. Together, these results suggest that users of `XomicsToModel` can expect to obtain thermodynamically consistent, biologically faithful models that are immediately applicable to hypothesis generation and disease-oriented research.

## ACKNOWLEDGEMENTS

The authors would like to thank Hanneke Leegwater for her helpful feedback while revising the manuscript.

## AUTHOR CONTRIBUTIONS

CRediT (Contributor Roles Taxonomy) author statement

German Preciat: Conceptualisation, Methodology, Software, Tutorial and Writing - Original draft; Agnieszka Wegrzyn: Methodology, Software, Data Curation and Writing - review and editing; Xi Luo: Methodology and Software; Ines Thiele: Conceptualisation, Methodology, Software and Funding acquisition; Thomas Hankemeier: Resources, Supervision and Funding acquisition, and Ronan M.T. Fleming: Conceptualisation, Methodology, Software, Writing - Original Draft, Writing - review and editing, Supervision and Funding acquisition

## DATA AND CODE AVAILABILITY

Constraint-based reconstruction, modelling and analysis was implemented in MATLAB (MathWorks Inc.). Open source computer code enabling the reproduction of all computational steps is available within the COBRA Toolbox [38] version 3.4+. This includes code for model generation `XomicsToModel.m`, thermodynamically feasible model extraction `thermoKernel.m`, debugging the performance of `XomicsToModel debugXomicsToModel.m`, comparison of multiple models `compareXomicsToModel.m`, and for maximisation of flux entropy, with the option of quadratic penalisation of deviation from measured experimental fluxes `entropicFBA.m`. See: https://github.com/opencobra/cobratoolbox/tree/master/src/dataIntegration/XomicsToModel

An executable narrative tutorial demonstrating the use of the XomicsToModel pipleine is also available `tutorial_XomicsToModel.mlx`, with the example data for generating the a dopaminergic neuronal metabolic model.

See https://github.com/opencobra/COBRA.tutorials/tree/master/dataIntegration/XomicsToModel

A html version of this tutorial is accessible here: https://opencobra.github.io/cobratoolbox/stable/tutorials/tutorialXomicsToModel.html

Linear and quadratic optimisation problems may be solved with open source solvers, e.g., GLPK, or using an industrial solver, such as Gurobi 9.1 (Gurobi Inc). Nonlinear convex optimization problems, e.g., entropic optimisation, may be solved with the open source solver *Primal-Dual interior method for Convex Objectives* PDCO implemented in MATLAB (MathWorks Inc.), or with an industrial solver, using the exponential cone solver within Mosek 10.0.30 (Mosek ApS). Note that PDCO may require specialist numerical optimisation parameters to be tailored to each model while Mosek is more robust with respect to the numerical characteristics of input models. Each of the aforementioned solvers is interfaced with the `COBRA Toolbox` and are suited for models are not numerically ill-scaled.

